# Subcutaneous delivery of an antibody against SARS-CoV-2 from a supramolecular hydrogel depot

**DOI:** 10.1101/2022.05.24.493347

**Authors:** Catherine M. Kasse, Anthony C. Yu, Abigail E. Powell, Gillie A. Roth, Celine S. Liong, Carolyn K. Jons, Awua Buahin, Caitlin L. Maikawa, Sawsan Youssef, Jacob E. Glanville, Eric A. Appel

## Abstract

Prolonged maintenance of therapeutically-relevant levels of broadly neutralizing antibodies (bnAbs) is necessary to enable passive immunization against infectious disease. Unfortunately, protection only lasts for as long as these bnAbs remain present at a sufficiently high concentration in the body. Poor pharmacokinetics and burdensome administration are two challenges that need to be addressed in order to make pre- and post-exposure prophylaxis with bnAbs feasible and effective. In this work, we develop a supramolecular hydrogel as an injectable, subcutaneous depot to encapsulate and deliver antibody drug cargo. This polymer-nanoparticle (PNP) hydrogel exhibits shear-thinning and self-healing properties that are required for an injectable drug delivery vehicle. In vitro drug release assays and diffusion measurements indicate that the PNP hydrogels prevent burst release and slow the release of encapsulated antibodies. Delivery of bnAbs against SARS-CoV-2 from PNP hydrogels is compared to standard routes of administration in a preclinical mouse model. We develop a multi-compartment model to understand the ability of these subcutaneous depot materials to modulate the pharmacokinetics of released antibodies; the model is extrapolated to explore the requirements needed for novel materials to successfully deliver relevant antibody therapeutics with different pharmacokinetic characteristics.

## 1 Introduction

The COVID-19 pandemic has necessitated the accelerated development of a wide range of treatment and prevention strategies in addition to the rapid pursuit of vaccines. One strategy is the development of new broadly neutralizing antibody (bnAb) cocktails, which have been implemented clinically to treat severe disease and are now being investigated for use as pre- and post-exposure prophylaxis.^1,2^ Alongside the development of vaccines, passive immunization, or the direct administration of neutralizing antibodies to prevent disease, is an important tool to fight infectious disease. Passive immunization is a useful strategy against pathogens for which development of typical active vaccines is challenging due to high mutation rates and uncertain correlates of immunity, such as HIV, malaria, and Zika, among others.^3–7^ Once broadly neutralizing antibodies can be identified, passive immunization becomes a possibility. Protection from passive immunization is nearly instantaneous as it does not rely on stimulating the immune system to produce antibodies against a pathogen, a process which often takes weeks to achieve maximum protection.^8^ Circumventing the immune system also allows immune compromised individuals to be protected by passively transferred antibodies, rather than relying on herd immunity for protection.

Unfortunately, passively-transferred immunity is limited by the circulation lifetime of bnAbs, which is typically on the order of weeks when administered in the typical manner by intravenous infusion or subcutaneous bolus injection (Figure 1). ^9^ This short duration of protection necessitates repeated administration, which leads to increased burden, poor patient adherence, and limited global reach of this strategy. In addition, most monoclonal antibody formulations are delivered intravenously, which limits the ability to quickly deploy doses to patients as IV infusions require more time and usually take place in a hospital or clinic setting.^10^ Thus, despite its potential, passive immunization is generally impractical with the poor pharmacokinetics exhibited by current formulations. A solution would be to increase bnAb circulation time, with the goal of a single injection conferring protection for upwards of six months. Common approaches to increasing the circulation time include injecting subcutaneously and leveraging recent advances in antibody modification techniques to enhance circulation half-lives.^11,12^ With the advent of antibody engineering techniques, one successful approach to increase circulation half-life in second-generation bnAbs is to introduce modifications into the crystallizable fragment (Fc) domains, which have been shown to nearly triple half-life.^11,13^ The development of bnAbs in general has largely been advanced by researchers pursuing passive prophylaxis against HIV, and the first long half-life bnAbs with Fc modifications are currently undergoing clinical trials.^14,15^

**Figure 1:**
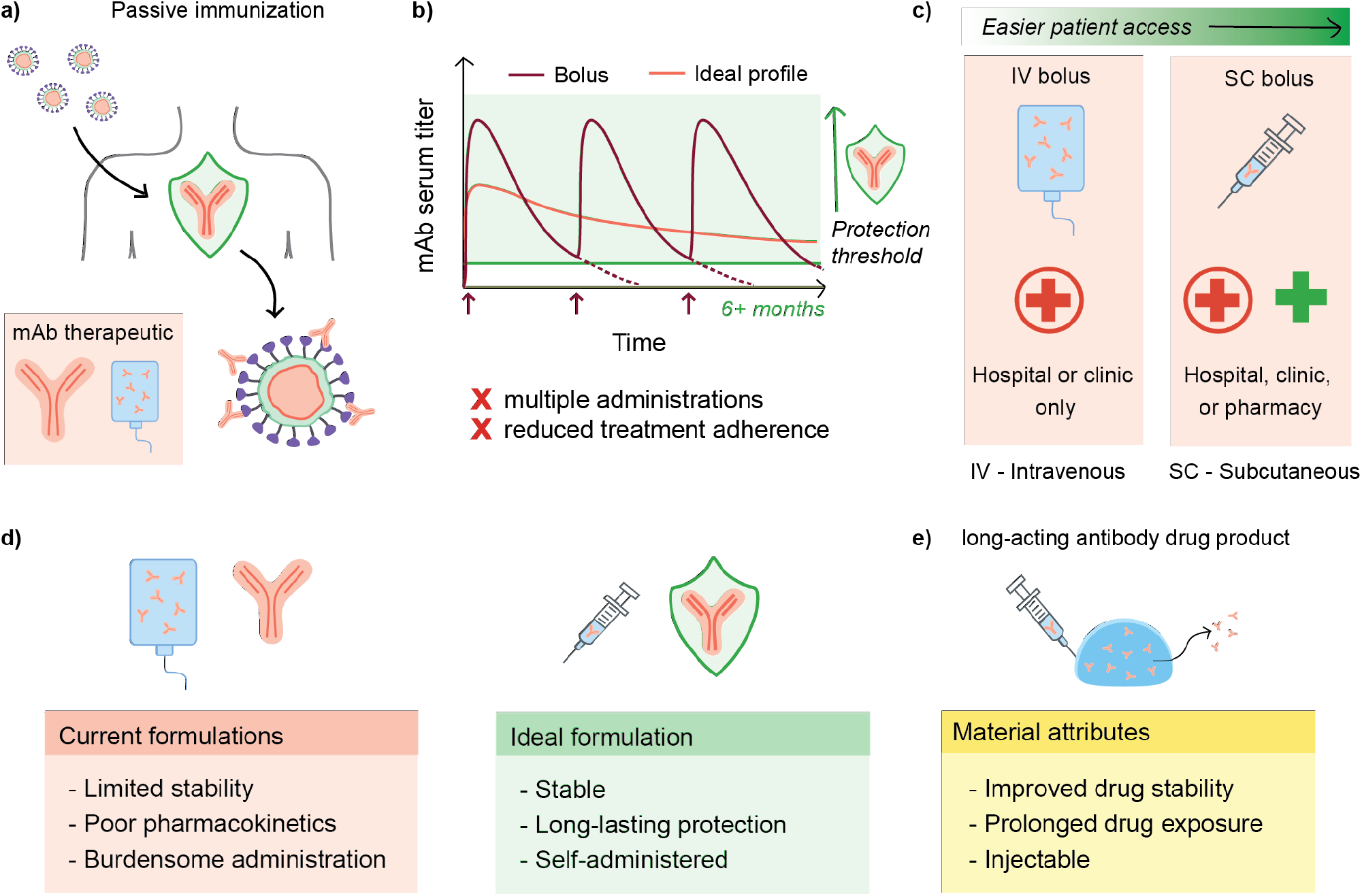
Passive immunization enabled by delivery from a subcutaneous depot. a) Passive immunization provides protection against infectious disease through direct administration of broadly neutralizing antibodies (bnAbs). b) Current bnAbs exhibit poor pharmacokinetics and require multiple doses to maintain a protective titer; in addition, c) most bnAbs are administered by IV infusion, which limits the feasibility of the treatment. d) Comparing current formulation limitations with those of an ideal formulation, e) a subcutaneous depot containing bnAbs emerges as a potential solution to current formulation challenges.

An alternative and complementary method towards improving circulation time of bnAbs is to develop new materials capable of acting as long-term depots of bnAbs and providing optimal pharmacokinetic profiles. Antibodies share many of the same delivery challenges as other protein biotherapeutics, such as being prone to aggregation and instability in formulation.^10,16^ A variety of injectable materials for controlled delivery of biotherapeutics have been explored.^10,17,18^ Recent materials used for encapsulation and delivery of antibodies include alginate hydrogels, PLGA nanoparticles, peptide hydrogels, agarose-desthiobiotin/streptavidin-antibody hydrogels, silk hydrogels, and other materials of different architectures.^19–26^ However, although many systems have been explored, a few critical materials challenges have limited their translation to the clinic. For example, materials platforms relying on in situ gelation or strong ionic crosslinking often exhibit significant burst release of encapsulated drug cargo when formulated to allow for injectability. Moreover, the exact materials and kinetic parameters required to achieve long-term pharmacokinetics is difficult to delineate due to the complex relation between the bnAbs, materials architecture, and biological functions in vivo.

An ideal antibody formulation will be stable both on the shelf and in vivo, can be administered outside of a clinic setting (e.g. subcutaneous injection rather than IV), and will exhibit appropriate pharmacokinetics to provide long-term protection to reduce patient burden and increase compliance. Given these desired attributes, we hypothesized that our supramolecular polymer-nanoparticle (PNP) hydrogels would be an appropriate vehicle to encapsulate and deliver bnAbs from a subcutaneously injected depot to result in more optimal pharmacokinetic profiles. These PNP hydrogels have been shown to be injectable, biocompatible in vivo, and drug-stabilizing and have been shown to be effective for a range of biomedical applications including cell encapsulation and delivery, vaccine delivery, cancer immunotherapy, and adhesion prevention. ^27–41^ In this work,we study the suitability of the PNP hydrogel platform to effectively deliver antibodies against SARS-CoV-2, and more broadly, we seek to investigate what is required to effectively design materials to act as a subcutaneous delivery depot for antibody therapeutics. We first evaluate the rheological characteristics, in vitro antibody release kinetics, and antibody stability in the PNP hydrogel system. Then, a selected candidate PNP hydrogel formulation is evaluated in vivo as a subcutaneous drug delivery depot for a novel neutralizing antibody against SARS-CoV-2. Finally, we use compartment modeling to develop our understanding of how a subcutaneous depot can alter drug pharmacokinetics and inform the design process for next-generation biomaterials.

## 2 Materials and Methods

### 2.1 Materials

Dodecyl-modified (hydroxypropyl)methyl cellulose (HPMC-C_12_, made from USP-grade HPMC (hypromellose, Sigma-Aldrich)) and poly(ethylene glycol)-block-poly(lactic acid) (PEG-b-PLA) block copolymers were synthesized as previously described; PEG-b-PLA nanoparticles (NPs) were assembled by nanoprecipitation as previously described.^27,28,42^ Unless otherwise stated, all chemicals were obtained from Sigma-Aldrich. Purified rat IgG was obtained from MP Biomedicals as a lyophilized powder. The Centi-C10 antibody against SARS-CoV-2 was generously provided by Dr. Sawsan Youssef and Dr. Jake Glanville at Centivax.

### 2.2 PNP hydrogel formulation

PNP formulation nomenclature is denoted by network polymer wt% : nanoparticle wt% (P:NP). For example, a 2:10 PNP hydrogel formulation contains 2 wt% HPMC-C_12_, 10 wt% self-assembled PEG-PLA nanoparticles, and 88% buffer. Stock solutions of the two individual components were prepared at 6 wt% HPMC-C_12_ and 20 wt% NPs, both in phosphate buffered saline (PBS). Supramolecular hydrogels were formed by mixing the stock solutions of the two components with additional buffer as needed to achieve the desired final component concentrations. For formulations containing antibodies, the drug cargo was incorporated at the desired concentration in the buffer component. For in vitro release assays and diffusion measurements, a total concentration of 10 mg/mL rat IgG was loaded into the hydrogels (9 mg/mL rat IgG and 1 mg/mL fluorescently-labeled rat IgG).

### 2.3 Dynamic and flow rheometry

All rheometry experiments were performed on a torque-controlled Discovery Hybrid Rheometer (DHR2, TA Instruments). Frequency sweeps were performed at 0.5% strain from 0.01 to 10 s^-1^ (in the linear viscoelastic regime of these materials) on a 20 mm parallel plate geometry. Flow sweeps were performed from high to low shear rates on a 40 mm 2° cone geometry. Step shear and stress-controlled yield stress experiments were performed using a 20 mm serrated parallel plate geometry. Prior to the step shear experiment, samples were first pre-conditioned for 60 s at 0.1 rad/s followed by 30 seconds at 10 rad/s. Step shear experiments were performed with a peak hold at 0.1 rad/s for 180 s followed by 30 s at 10 rad/s, repeated for four cycles. The viscosity recovery curves for the last three cycles were fit to the equation *y* = *A* * (1 – *e*^-x/*τ*^) +*B*, where *τ* is the characteristic recovery time. Stress controlled yield stress measurements were performed from low to high stress with steady state sensing and 10 points per decade.

### 2.4 Fluorescently labeled HPMC-C_12_ and rat IgG

HPMC-C_12_ was made as described, except after precipitation from acetone, the polymer was dried under high vacuum and then re-dissolved in 40 mL of *N*-methyl-2-pyrrolidone (NMP). 10 mg of fluorescein isothiocyanate isomer I (FITC) was dissolved in 10 mL of NMP and added dropwise to the reaction flask. 3-4 drops of N,N-Diisopropylethylamine (DIPEA) was added and the reaction was allowed to proceed for 16 hrs. The labeled polymer was then purified according to the same methods cited above for synthesis of unlabeled HPMC-C_12_. Fluorescently labeled rat IgG was prepared by adding 16 μL of 2 mg/mL FITC in PBS to 3 mg of rat IgG in 1 mL of PBS and incubating at 4 °C for 24 hrs. The solution was then passed through a PD-10 MidiTrap desalting column to remove free FITC, lyophilized, and then re-dissolved at 60 mg/mL in PBS as a stock solution.

### 2.5 In vitro release assays

Burst release assays were conducted by injecting 100 μL of PNP hydrogel loaded with fluorescently-labeled rat IgG into a microcentrifuge tube filled with 400 μL of PBS. After 5 minutes, 300 μL of the supernatant was removed and analyzed to determine the amount of FITC-labeled IgG using a Synergy H1 Hybrid Multi-Mode Plate Reader (BioTek). Release assays were conducted by adding 100 μL hydrogel to microcentrifuge tubes and then centrifuging until the hydrogel was situated in the bottom of the tube. Then PBS was added on top of the hydrogel. At regular intervals, aliquots of the supernatant were taken for measurement in the plate reader as described above, and the sampled volume was replaced with PBS.

### 2.6 In vitro diffusivity measurements

All fluorescence recovery after photobleaching (FRAP) experiments were conducted in the Stanford University Cell Sciences Imaging Facility (CSIF) at room temperature or 37 °C. FRAP experiments were performed on PNP hydrogels loaded with fluorescently labeled rat IgG, or on PNP hydrogels made with fluorescently lableled HPMC-C_12_. An inverted Zeiss LSM 780 Laser Scanning Confocal Microscope (Germany) with a Plan-Apochromat 20x/0.8 M27 objective lens was used with ZEN lite software (Zeiss). A 488 nm argon laser at 20% intensity was used to excite the fluorescein for imaging. 405 and 488 nm argon lasers set at 100% intensity were used for photobleaching. A high-voltage limit was set at 700 V to reduce noise. The samples were placed on sterile 0.18 mm thick glass bottom *μ*-dish (Ibidi). Each batch of PNP hydrogel was split into 3 different samples in the μ-dish and measurements were performed 2-3 times on each sample. For each test, 10 pre-bleach images were taken, followed by photobleaching of a 25 nm diameter circle for 10 cycles at a pixel dwell time of 177.32 s. At a minimum, 500 post-bleach frames were recorded to form the recovery curve, but up to 15 min of fluorescence recovery data was recorded for slower recovery samples. Data was corrected for fluorescence signal drift due to photobleaching. The diffusion coefficient was calculated as 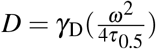 where γ_D_ is a constant that depends on beam shape, transport type, and bleaching conditions, *ω* is the radius of the bleached area, and τ_0.5_ is the time to recover 50% of the initial measured fluorescence intensity.^43^ Diffusivity of freely diffusing rat IgG was determined using dynamic light scattering (DLS, n=5).

### 2.7 Centi-C10 generation

Centi-C10 is a high affinity monoclonal antibody (human-IgG1) that binds the wild type SARS-CoV-2 Receptor binding domain (RBD). This clone blocks the interaction between the RBD and human-ACE-2. Centi-C10 was isolated from an ScFV phage library constructed using the parental antibody 6W41(CR3022-originally binds SARS-CoV-1). The library were built according to the proprietary protocol of affinity maturation /humanization library build (Centivax Inc./Distributed Bio/ Charles River Laboratories). Briefly, the 6 CDRs of the original clone were synthesized with scanned mutations by overlapping PCR reactions to engraft it into the appropriate human germlines used by the company. The libraries then were transformed to around a billion copies per library. The library was panned using the wild type-RBD-Fc-Avi tag construct generated by Centivax Inc. The library was subjected to 4 rounds of selection and ScFv were tested for the binding of SARS-CoV-2 RBD and blocking its interaction with of hu-ACE-2 on high-throughput surface plasmon resonance (SPR) on Carterra LSA Array SPR instrument (Carterra) equipped with HC200M sensor chip (Carterra) at 25 °C.

### 2.8 Antibody cloning into expression vectors for transient transfection

Heavy chain and light chain sequences of the complimentary determinant regions (CDRs) of centi-C10 ScFv was cloned into each constant region of human IgG1 heavy chain, and human kappa constant IGKC, in the mammalian expression pTT5 vector (Licensed from National Research Council of Canada, Toronto Canada), respectively. Centi-C10 was generated in Transient Transfection using human embryonic kidney Expi293F^™^ ThermoFisher Scientific (San Diego, California), according to manufacture protocol at Centivax Inc. (South San Francisco). The antibody was then purified using HiTrap MabSelect PrismA protein A column (Cytiva, Marlborough, MA,), pre-equilibrated with at least 5 column volume (CV) of 1x PBS, pH 7.9 (ThermoFisher Scientific), in AKTA pure 25M fast protein liquid chromatography (FPLC) system (Cytiva). Flow rate was adjusted accordingly depending on the protein titer to achieve appropriate dynamic binding capacity.

### 2.9 Spike-pseudotyped lentivirus production and viral neutralization assays

Spike-pseudotyped lentivirus was produced in HEK293T cells as previously described. ^44^ Briefly, HEK293T cells cultured in D10 medium (DMEM + additives: 10% fetal bovine serum, L-glutamate, penicillin, streptomycin, and 10 mM HEPES) were seeded at a density of 6 million cells in 10-cm dishes 1 day prior to transfection. Cells were transfected using calcium phosphate transfection with 5 plasmids previously described in^45^ in the following ratios: 10 *μ*g luciferase-containing lentivirus packaging vector (pHAGE-Luc), 3.4 *μ*g FL SARS-CoV-2 Spike, 2.2 *μ*g HDM-Tat1b, 2.2 *μ*g pRC-CMV-Rev1b, and 2.2 *μ*g HDM-Hgpm2. Culture medium was exchanged ~18-24 hrs post transfection and virus was harvested from cell supernatant ~72 hrs post transfection by centrifugation at 300 x g for 5 min followed by filtration through a 0.45 *μ*m syringe filter. Viral stocks were aliquoted and stored at −80 °C. Spike-pseudotyped lentiviral neutralization assays were performed using HeLa cells overexpressing human ACE2, as described in. ^46^ HeLa/ACE2 cells were plated in 96-well clear bottom, white-walled plates 1 day prior to infection at a density of 5,000 cells per well. Mouse serum was heat inactivated for 30 min at 56 °C and diluted in D10 medium; dilution was 1:50 for measuring neutralization at day 7. Virus was diluted with D10 medium and supplemented with polybrene (present at 5 *μ*g/mL final concentration in assay wells). Diluted mouse serum and virus were incubated at 37 °C for 1 hr. D10 media was aspirated off of plated cells and replaced with virus/serum dilutions for ~48 hrs at 37 °C. Viral infectivity was read out using BriteLite (Perkin Elmer) luciferase readout solution and relative luminescence units (RLUs) were quantified using a BioTek plate reader. All samples assayed in duplicate.

### 2.10 In vitro stability assay

Centi-C10 samples were prepared at 1 mg/mL in either PBS buffer or encapsulated in a 2:10 PNP hydrogel as described above. Samples were loaded into 7 mL glass scintillation vials at a volume of 0.5 mL each. The vials were sealed, wrapped in Parafilm™, and affixed on their sides (to maximize air-water interface area and accelerate degradation) to the platform of a rotator plate inside an incubator. Samples were incubated at 37 °C and constantly agitated at 200 rpm. Aliquots taken at weekly timepoints up to 4 weeks were analyzed via ELISA.

### 2.11 Centi-C10 ELISA

Centi-C10 concentrations were measured by an anti-RBD (receptor binding domain) ELISA. SARS-CoV-2 (2019-nCoV) Spike RBD-His Recombinant Protein (SinoBiological, 40592-V08H) at 2 *μ*g/mL in PBS was coated onto 96-well clear, round bottom MaxiSorp plates (Thermo Scientific) by overnight incubation at 4 °C. All reagents were added to each well in 50 *μ*L quantities unless otherwise stated. All incubation steps were at room temperature. Plates were washed 5x with 250-300 *μ*L PBS-T (0.05% Tween 20) and subsequently blocked for 1 hr with 250 *μ*L of 1% bovine serum albumin (BSA) in PBS. Samples (either in vitro or mouse serum) were prepared by serial dilution in 1% BSA. Plates were washed 5x in PBS-T before adding samples and incubating for 2 hrs. Plates were again washed 5x in PBS-T before adding a 1:10,000 dilution in 1% BSA of the secondary antibody, Peroxidase AffiniPure F(ab’)_2_ Fragment Goat Anti-Human IgG, Fcγ fragment specific (Jackson Immunoresearch, 109-036-008, RRID: AB_2337591), and incubating for 1 hr. Plates were washed 5× in PBS-T before developing with TMB ELISA Substrate (High Sensitivity) (Abcam, ab171523). Plates were allowed to develop for 3 min before the reaction was stopped by adding 1 N HCl. Absorbance at 450 nm was measured with a Synergy H1 Hybrid Multi-Mode Plate Reader (BioTek). Every plate contained a 16 point Centi-C10 standard curve assayed in duplicate. The standard curve was fit using a four-parameter doseresponse curve (variable slope) in GraphPad Prism 9. To quantify the concentration of individual samples from their respective serial dilution series, the dilution with an absorbance value nearest to that of the EC50 of the standard curve was interpolated.

### 2.12 Pharmacokinetic (PK) modeling

The IV bolus data was fit to a two-phase exponential decay with the plateau constrained to zero in GraphPad Prism 9 to determine *k_elim_; V_d_* was calculated from M_0_/C_0_, where C_0_ is the initial concentration from the two-phase exponential. The SC bolus data was fit to a single compartment pharmacokinetic model. ^47^ Simulated pharmacokinetic profiles were generated in MATLAB. The differential equations and analytical solutions to the single compartment model and two compartment model incorporating a subcutaneous depot can be found in the supporting information.

### 2.13 In vivo Centi-C10 mAb pharmacokinetic study

All animal studies were performed in accordance with National Institutes of Health guidelines and with the approval of the Stanford Administrative Panel on Laboratory Animal Care (APLAC-32109). Female B6.Cg-Fcgrt^tm1Dcr^ Prkdc^scid^ Tg(FCGRT)32Dcr/DcrJ (The Jackson Laboratory, Stock No. 018441) mice age 12-14 weeks were administered Centi-C10 antibody via IV (retro-orbital) or SC injection under brief isoflurane anesthesia. All formulations were prepared at a concentration of 3.3 mg/mL Centi-C10 in PBS and delivered in 150 *μ*L volume (dose: 500 *μ*g/mouse). Hydrogel samples were prepared for the in vivo study from a stock solution of 4 wt% HPMC-C_12_ dissolved in PBS containing the appropriate concentration of Centi-C10 antibody and 20 wt% NPs stock solution. These samples were mixed by filling two syringes with HPMC-C_12_+ Centi-C10 stock solution and NP stock solution, respectively, and then attaching them to a luer lock elbow connector to push the components through for mixing. This process allows for mild encapsulation of the antibody and results in pre-loaded syringes for injection. Hydrogel components for in vivo studies were tested for bacterial endotoxins using a limulus amebocyte lysate (LAL) test (PYROSTAR ES-F, FUJIFILM Wako Chemicals) prior to use. Serum samples were collected via the tail vein at regular timepoints for analysis of Centi-C10 concentration by ELISA. For the IV and SC bolus groups, samples were collected at 1, 4, 8, 12, and 20 hrs; these timepoints were split across each group (n = 3). For the IV bolus, SC bolus, and SC gel groups, timepoints were taken at 24 hrs, and then 4, 7, 10, and 14 days followed by weekly timepoints until the end of the study (n = 6).

### 2.14 Statistical analysis

All statistical analyses were performed using GraphPad Prism 9. Results generally reported as mean ± SD unless specified otherwise. In general, comparison of two means was determined with a two-sided, unpaired t-test (τ, burst release, bioavailability). Diffusivity measurements and viral neutralization data were compared using a one-way ANOVA. Tukey post-hoc tests were applied to account for multiple comparisons (α = 0.05); adjusted p-values were reported.

## 3 Results and discussion

### 3.1 Rheological characteristics of the PNP hydrogel

PNP hydrogels are non-covalently crosslinked hydrogels formed through dynamic, multivalent interactions between the network polymer, hydrophobically modified hydroxypropyl methyl cellulose (HPMC-C_12_) and self-assembled polymeric nanoparticles formed through nanoprecipitation of polyethylene glycol-block-polylactic acid (PEG-b-PLA; d_h_ ≈ 30 nm) (Figure 2a). The hydrogel is formed by simply mixing aqueous solutions of the two components. The mild gelation conditions are conducive to encapsulating biotherapeutic drug cargo, such as antibodies, into the bulk material. PNP hydrogels can exhibit a range of rheological properties depending on the ratio of polymer to nanoparticles in the formulation and overall solid polymer content. A 2:10 PNP formulation indicates 2 wt% HPMC-C_12_ and 10 wt% PEG-PLA nanoparticles, with the remaining 88 wt% buffer plus drug cargo. Two formulations, 1:5 and 2:10, were selected for comparison. Both formulations exhibit solid-like behavior (G’> G’’, tan(δ) = 0.1-0.5) across the entire range of the frequency sweep shown in the linear viscoelastic regime (Figure 2b). The 2:10 formulation has a shear storage modulus approximately an order of magnitude greater than the 1:5 formulation.

**Figure 2:**
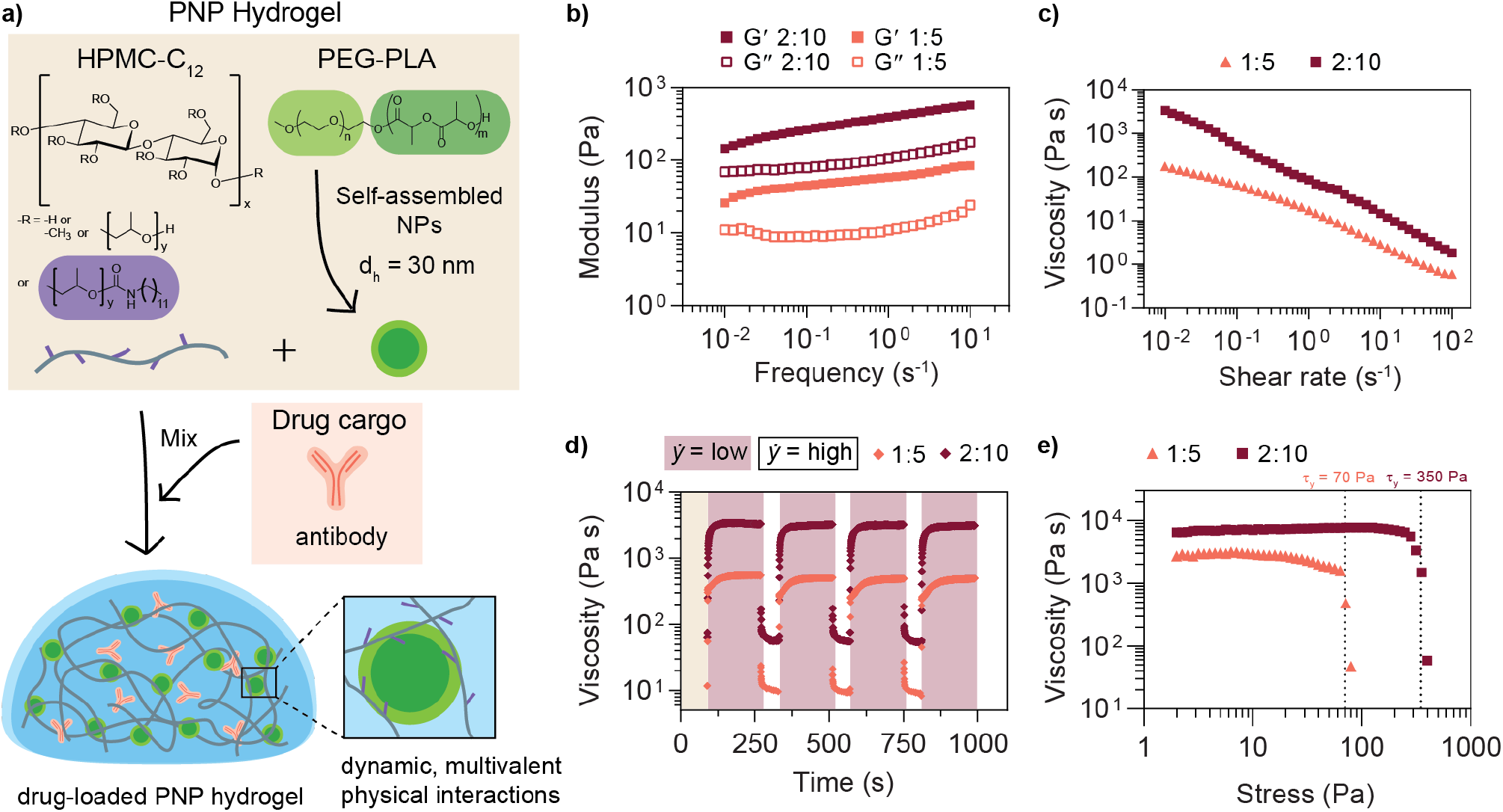
Polymer nanoparticle (PNP) hydrogel formulation and rheological properties. a) PNP hydrogels are dynamic networks formed from supramolecular interactions between a dodecyl-modified (hydroxypropyl)methyl cellulose (HPMC-C12) network and self-assembled nanoparticles composed of poly(ethylene glycol)-*block*-poly(lactic acid) (PEG-PLA) block copolymers. Drug cargo can be easily loaded into the hydrogel by incorporation into the buffer. Formulation nomenclature is denoted by network polymer wt% : nanoparticle wt%. b) Storage (G’) and loss (G’’) moduli and c) viscosity as a function of shear rate were measured for two different PNP formulations (data adapted from Meis, et al. ^27^). d) Step shear tests were performed by alternating between low (0.1 rad/s) and high (10 rad/s) shear rates after preconditioning for 90 seconds. e) Yield stresses (indicated by dotted lines) were determined from a stress-controlled sweep.

One way to assess injectability, a critical attribute for a subcutaneous delivery vehicle, is to measure the shear rate dependence of the viscosity. Both formulations show a decrease in viscosity of nearly three orders of magnitude with increasing shear rate and are shear-thinning (Figure 2c). Shear rates achievable on a rheometer are too low to simulate injection through a needle, so it is also important to confirm injectability by testing with a relevant syringe/needle combination.^48^ These formulations are also injectable through a 21G needle.^42^ Step shear experiments were performed to simulate the step-wise change in shear rate during injection and determine the ability of the network to recover solid-like properties after being subjected to high shear rates(Figure 2d). From the viscosity recovery curves at low shear rate, the characteristic recovery time, *τ*, was determined to be 42±16 s and 10±1 s for the 1:5 and 2:10 hydrogel formulations, respectively (Figure S3). Previous findings show that a P:NP stoichiometric ratio between 0.1 and 1 (P:NP = 0.2 for both formulations) promotes bridging between components and thus gelation in this PNP hydrogel system; the 2:10 network likely recovers more quickly compared to the network 1:5 due to higher overall content of polymer and nanoparticles to form bridging interactions. ^49^

Finally, the yield stress of each formulation was determined to assess the ability of the depot to retain its shape under the forces present the site of injection; it has previously been demonstrated that yield stress is an important rheological property for subcutaneous depot formation and retention. ^34,50^ Yield stresses of 70 and 350 Pa were measured for the 1:5 and 2:10 formulations, respectively (Figure 2e). Again, previous work has shown that increasing nanoparticle content results in higher yield stress values. ^49^ Both of these formulations exhibit sufficiently high yield stress to form a persistent depot in the mouse subcutaneous space upon injection. ^50^ In addition, the hydrogel is rheologically stable for more than 5 weeks at room temperature (Figure S2).

### 3.2 In vitro antibody release and diffusivity from the PNP hydrogel

Next, we evaluated the drug delivery capabilities of the two formulations in vitro. Rat IgG was used as the model antibody in these in vitro experiments as most bnAbs are IgG type antibodies and generally similar in size and structure. We hypothesized that a faster recovery time, τ, in the step shear experiments would correlate with reduced burst release of encapsulated drug cargo as the network reforms more quickly upon injection (Figure 3a). To test burst release, hydrogels loaded with fluorescently labeled rat IgG were injected into microcentrifuge tubes filled with PBS and the amount of antibody released after 5 min was measured by fluorescence intensity. The 1:5 hydrogel released 16 ± 6% of the total cargo mass 5 minutes after injection, while the 2:10 hydrogel released only 1 ± 1% of the cargo; this result corroborates the hypothesis that materials exhibiting faster recovery times retain more cargo and reduce burst release (Figure 3b). Minimal burst release from the depot is critically important for long-term biotherapeutic delivery because the immediate loss of a substantial portion of the delivered dose can reduce the amount of time it is possible to maintain a protective titer. More broadly, burst release is concerning for therapeutics that exhibit dose-dependent toxicity; while bnAbs are generally well-tolerated even in high doses, this material could also be used for other applications where toxicity is a concern, such as in cancer immunotherapy. ^17,35^

**Figure 3:**
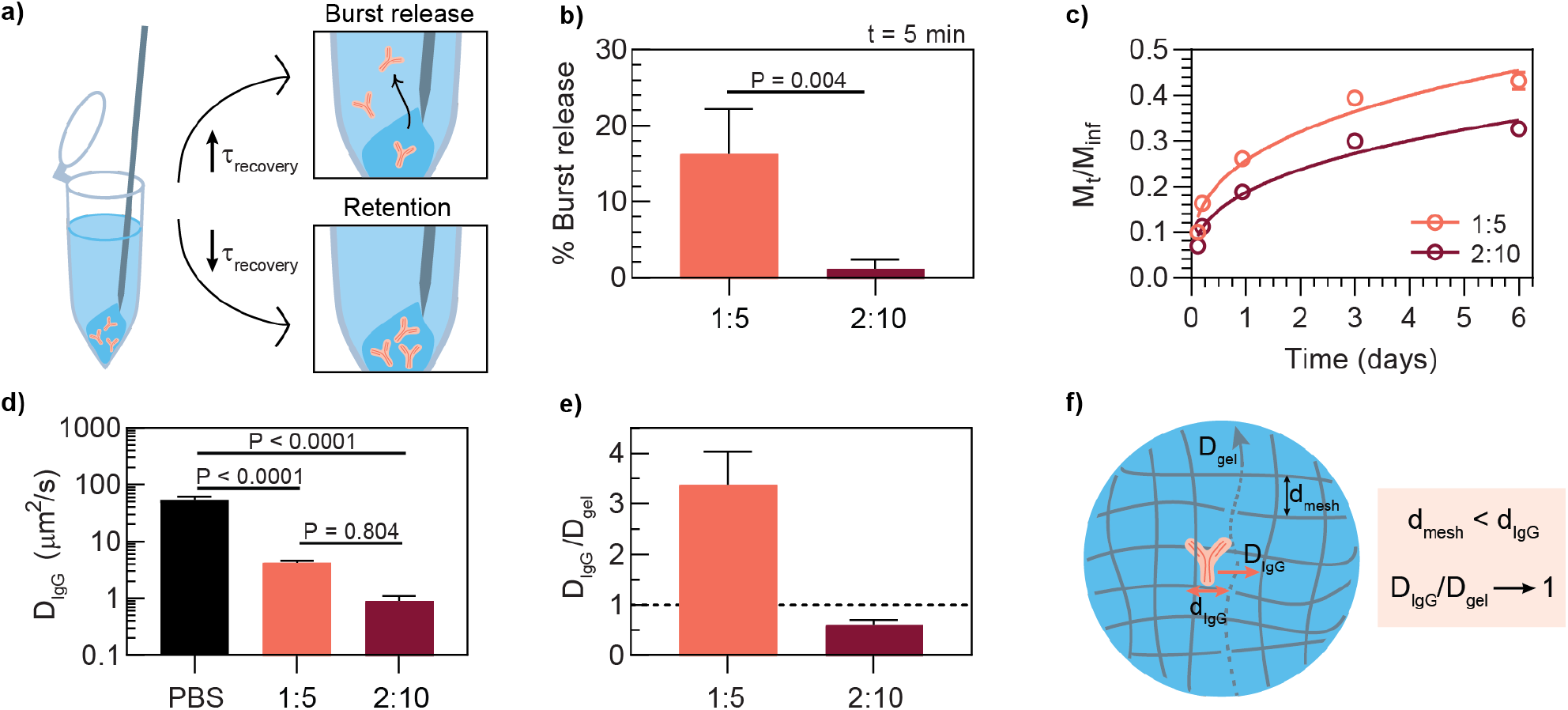
In vitro antibody release and diffusivity from the PNP hydrogel for two different hydrogel formulations. The selected antibody was fluorescently-labeled rat IgG. a) Characteristic recovery time *τ* correlates with burst release versus retention of antibody cargo. b) Percent of antibody released 5 minutes post-injection into a PBS-filled microcentrifuge tube (mean ± SD, n=3); p-value determined by two-sided unpaired t-test. c) Cumulative in vitro release of antibody from a PNP hydrogel (mean ± SD, n=5); trendline shown is a Ritger-Peppas empirical fit. ^51^ d) Diffusivity (DIgG) of free antibody compared to gel-encapsulated antibody as quantified by dynamic light scattering (DLS, PBS sample) and fluorescence recovery after photobleaching (FRAP, hydrogel samples) (mean ± SD, n=5 for PBS, n=3 for hydrogels). Statistical significance was determined by one-way ANOVA (F (DFn, DFd) = F (2, 8) = 82.62). Tukey post-hoc tests were applied to account for multiple comparisons. e) Ratio of antibody diffusivity to hydrogel diffusivity (D_gel_), where D_gel_ is the diffusivity of fluorescently-labeled HPMC-C_12_ within the denoted hydrogel formulation as measured by FRAP (mean ± SD, n=3). f) Illustration of possible antibody diffusion process where passive diffusion of the antibody is limited by the dynamic mesh of the hydrogel network.

Drawing from the rheological characteristics, increasing storage modulus typically correlates with reducing the mesh size of the hydrogel network; therefore, we hypothesized that the hydrogel formulation with greater storage modulus would be more effective at slowing antibody release. The sustained release capabilities of the two PNP formulations were evaluated by measuring the amount of fluorescent rat IgG released into buffer over time from hydrogels in microcentrifuge tubes. The higher modulus 2:10 hydrogel released a smaller percentage of the total mass (M_t_/M_inf_ = mass at time, t / total mass) compared to the 1:5 formulation over 6 days (Figure 3c).

To develop a better understanding of how encapsulated drug cargo diffuses within the hydrogel, fluorescence recovery after photobleaching (FRAP) experiments were performed with fluorescently labeled rat IgG to determine D_IgG_, diffusivity of IgG, and fluorescently labeled HPMC-C_12_ to determine D_gel_, diffusivity of network polymer, in each formulation. Measured D_IgG_ in the 1:5 formulation (D_IgG_=4.3±0.4 *μ*m^2^/s) was about 4 fold greater than in the 2:10 formulation (D_IgG_=0.9±0.2 *μ*m^2^/s) (Figure 3d). These values are both an order of magnitude smaller than the diffusivity of freely diffusing IgG in PBS (D_PBS_=53.4 ± 9 *μ*m^2^/s), demonstrating that the PNP hydrogel constrains passive diffusion. An important observation is that D_IgG_ in the 2:10 formulation is nearly the same as the diffusivity of the HPMC-C12 chains themselves (D_gel_) (Figure 3e, Table S1). This data suggests that in the limit where the network mesh size is smaller than the size of the IgG, D_IgG_ is dictated by D_gel_, presenting a way to control D_IgG_ through optimization of the strength and kinetics of the supramolecular crosslinks and network polymer mobility in the case where D_IgG_/D_gel_ approaches 1. (Figure 3f). This data also suggests that the 1:5 hydrogel has a larger effective mesh size compared to the 2:10 hydrogel, allowing the IgG cargo to diffuse more quickly, whereas the 2:10 hydrogel mesh more effectively limits diffusion. Previous work indicates that the mesh radius of a 2:10 hydrogel network is approximately 6 nm; monomeric human IgG has a hydrodynamic radius of 5-6 nm, so it is likely to be entrapped by the hydrogel network. ^40,52,53^

Taken together, the rheological properties, in vitro release studies, and diffusion measurements indicate that the PNP platform, and particularly the 2:10 formulation, might be suitable for subcutaneous antibody delivery based on the desired attributes of injectability and prolonged drug exposure. The 2:10 formulation was selected for an in vivo antibody pharmacokinetics study based on its increased drug retention upon injection and slower cargo release over time compared to the 1:5 formulation. Lastly, we hypothesize that higher polymer content formulations (e.g. 3:15) would would not be beneficial because increasing the solid content in the formulation also generally reduces how much antibody drug can be encapsulated in the network, so higher polymer content formulations were not investigated for this work.

### 3.3 In vitro stability of an antibody against SARS-CoV-2

Centi-C10 is a high affinity monoclonal antibody (human-IgG1) that binds the wild type SARS-CoV-2 Receptor binding domain (RBD). This clone blocks the interaction between the RBD and human-ACE-2. The IC_50_ of Centi-C10 was determined to be 0.2 *μ*g/mL in a neutralization assay against spike-pseudotyped lentivirus (Figure 4).

**Figure 4:**
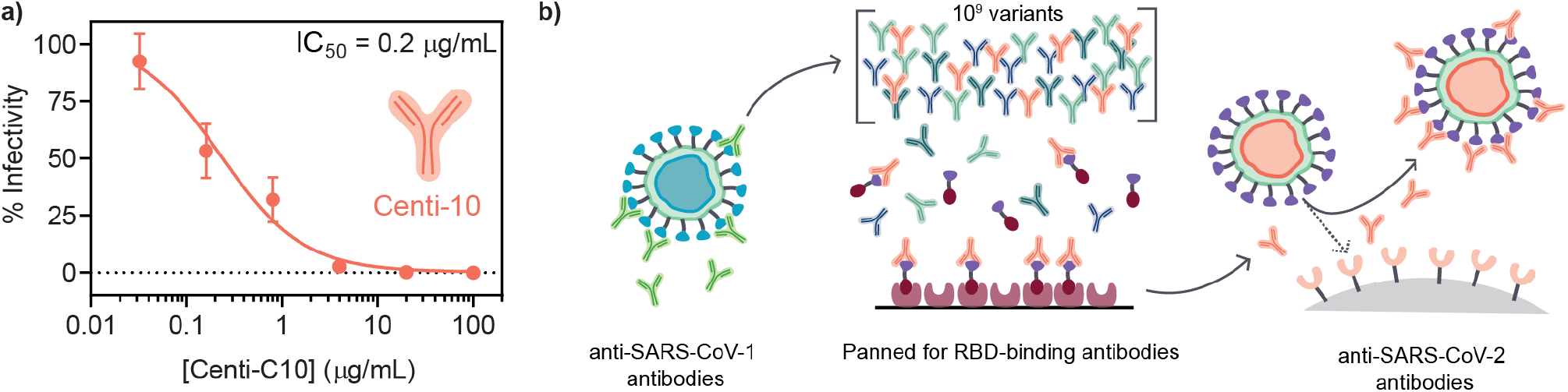
Centi-C10 antibody properties and discovery process. a) Centi-C10 potency quantified by viral neutralization assays of spike-pseudotyped lentivirus (mean ± SD, assayed in duplicate). b) Antibodies capable of binding the SARS-CoV-2 receptor binding domain (RBD), and thus blocking the binding of the virus to ACE-2 receptors on human cells, were discovered by multiple rounds of panning and selection from an ScFV phage library built from a SARS-CoV-1-binding parental antibody.

Prior to initiating an in vivo study, an in vitro stability assay was conducted to verify that the Centi-C10 antibody would be stable when encapsulated in the hydrogel. Centi-C10 samples prepared at 1 mg/mL in either PBS or 2:10 hydrogel were aged in glass vials at 37 °C with constant agitation on a shaker plate for up to 4 weeks. Weekly aliquots were analyzed via SARS-CoV-2 spike RBD ELISA as a measure of functional activity and stability (Figure 5a). After 1 week, the hydrogel-encapsulated Centi-C10 exhibited nearly unchanged binding activity in the assay compared to a fridge-stored control, while the PBS sample had only 37 ± 3% binding activity (Figure 5b). After 3 weeks, the Centi-C10 in PBS exhibited <10% binding activity whereas the encapsulated Centi-C10 still measured 40 ± 13%. Even after 1 week at 50 °C with constant agitation, hydrogel-encapsulated Centi-C10 exhibited nearly 60% binding activity (Figure S4). It is anticipated that both formulations would likely be more stable in vivo as the conditions of the in vitro assay were intended to accelerate degradation of the antibody. In addition, these assays not only indicate that antibodies should be stable within the hydrogel as a subcutaneous depot, but also that hydrogel-encapsulated antibody formulations could be more shelf-stable and cold-chain resilient. Shelf stability and cold-chain resilience are both key factors to both reducing overall cost and improving biotherapeutic drug access. ^54^–^56^

**Figure 5:**
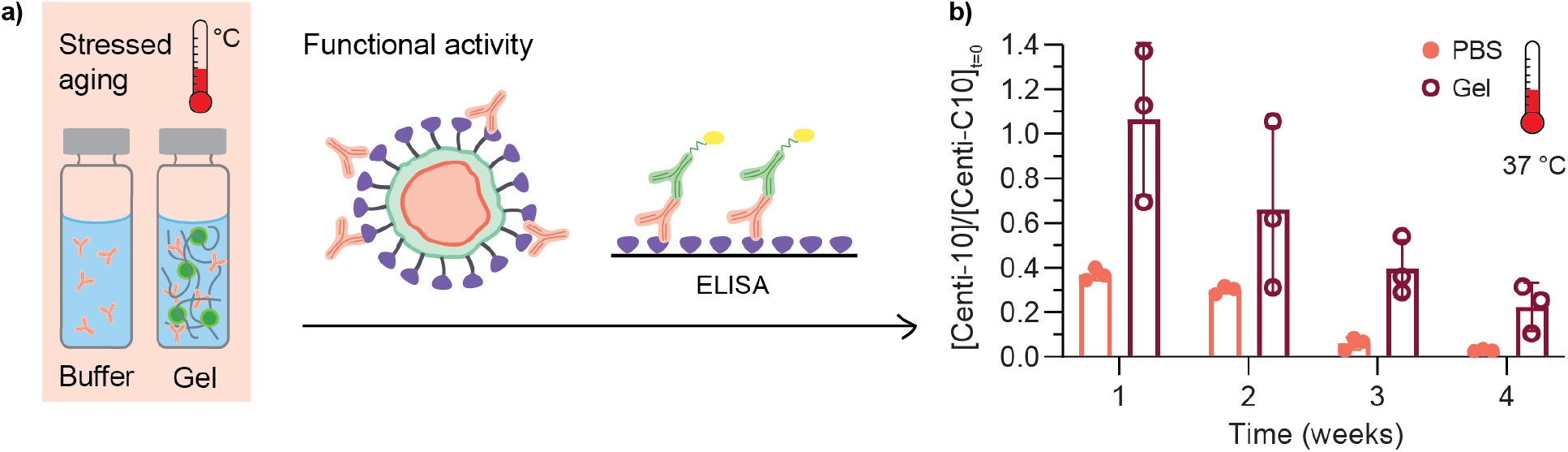
In vitro stability of Centi-C10 antibody in standard buffer formulation compared to PNP hydrogel-encapsulated Centi-C10. a) In vitro aging assays were conducted by subjecting samples in glass vials to heat and constant agitation. Aliquots were removed at weekly timepoints for analysis of functional stability by ELISA. b) Post-aging fraction of Centi-C10 binding to SARS-CoV-2 Spike RBD antigen by ELISA, normalized by a fridge-stored control (mean ± SD, assayed in triplicate).

### 3.4 Centi-C10 pharmacokinetics in a preclinical mouse model

To test whether the 2:10 hydrogel could serve as a subcutaneous antibody delivery depot, a study was conducted in *scid* FcRn-/-hFcRn (32) Tg mice where Centi-C10 pharmacokinetics were compared between standard bolus administrations and a subcutaneously injected hydrogel. These mice express an hFcRn transgene and are immunodeficient, allowing the long-term evaluation of human IgG antibody serum levels.^57,58^ Each mouse was administered a dose of 500 *μ*g of Centi-C10 via either IV bolus, SC bolus, or SC 2:10 hydrogel in PBS, and blood samples were collected throughout the time course for Centi-C10 serum concentration analysis via ELISA (Figure 6a).

**Figure 6:**
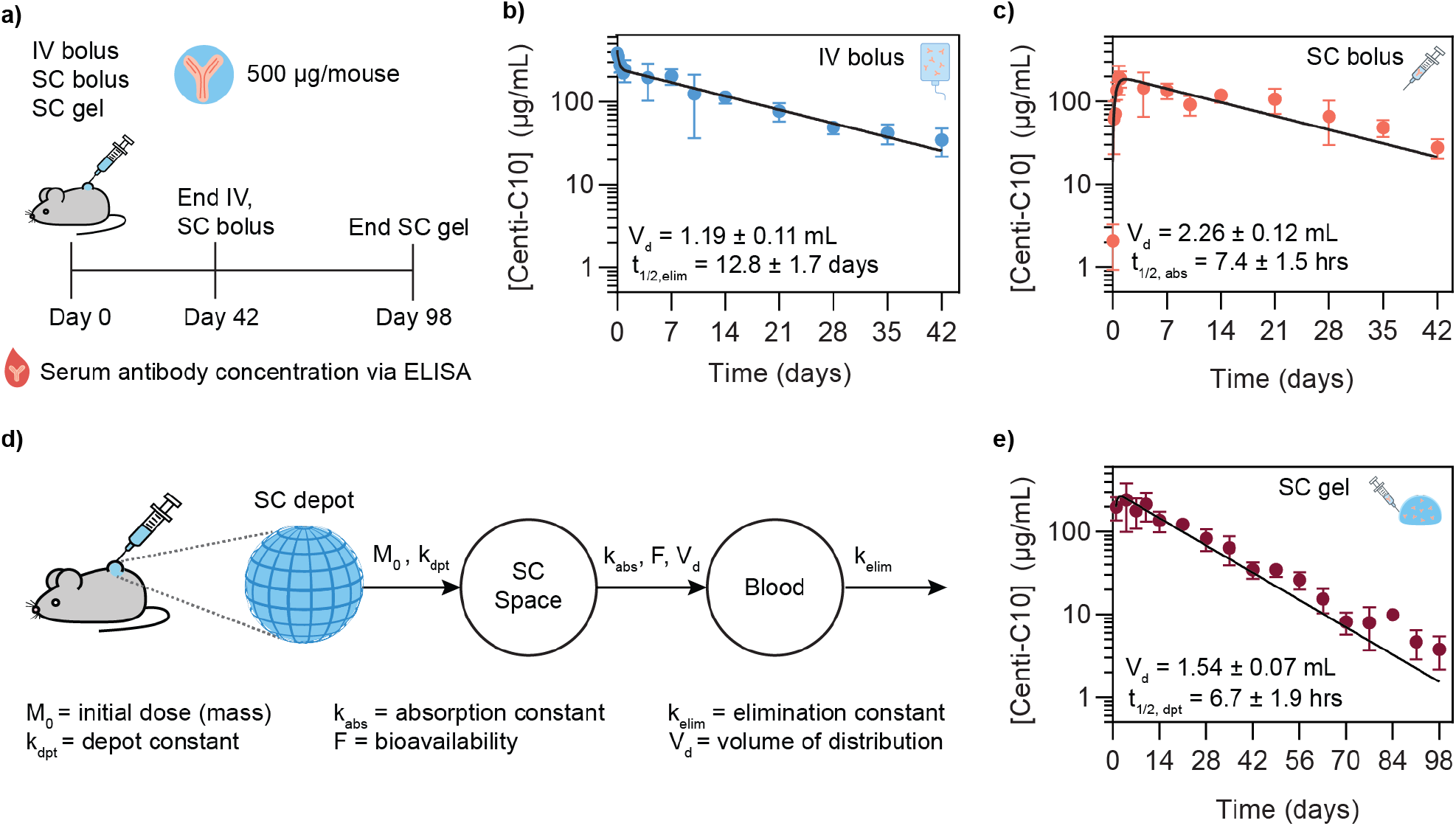
Centi-C10 pharmacokinetics in a preclinical mouse model. Centi-C10 antibody was administered to *scid* FcRn-/-hFcRn (32) Tg mice via three different routes and the pharmacokinetics were assessed as shown in a) via ELISA. Pharmacokinetic parameters were determined by fitting the Centi-C10 serum concentration profiles using a two-phase decay for the b) IV bolus or a single compartment model for the c) subcutaneous bolus. d) Two compartment model incorporating the subcutaneous depot that was applied to fit the pharmacokinetic profile of the e) SC gel group, using first order rate constants determined from the IV and SC bolus data. Data points shown as mean ± SD (n = 3 prior to 24 hrs, otherwise n = 6; for gel group, n ≥ 4 after day 63); calculated PK values shown as mean ± SE.

Antibody pharmacokinetics are often described using compartment modeling methods. ^47,59^ To understand how the subcutaneous hydrogel depot might modulate the serum pharmacokinetics, it was first necessary to quantify pharmacokinetic (PK) parameters for Centi-C10 in this mouse model from the IV and SC bolus data. The IV PK profile was fit with a two-phase exponential decay to yield a volume of distribution Vd,IV = 1.19 ± 0.11 mL and a serum elimination half-life t_1/2_,_elim_= 12.8 ± 1.7 days (Figure 6b). Next, the SC bolus data was fit with a standard one compartment PK model with first order rate constants, where t_1/2_,_elim_ was fixed to 12.8 days as determined from the IV bolus PK profile (Figure 6c). From this, the absorption half-life of Centi-C10 from the subcutaneous space was found to be t_1/2,abs_ = 7.4 ±1.5 hrs with a slightly larger V_d,SC_ = 2.26 ± 0.12 mL. These half-life values are similar to what might be expected from literature for IgG type antibodies delivered via IV or SC routes in this model. ^60,61^ Blood volume for a mouse is generally 1-2 mL depending on mass, so the V_d,IV_ is within the expected range. The Vd,SC from the one compartment model is nearly 2x greater than V_d,IV_, which is also not unexpected because antibodies delivered into the subcutaneous space are preferentially drained into the lymphatic system prior to entering circulation, so the drug should be distributed across more tissues in the body leading to an increase in V_d_.

To understand how the SC hydrogel depot might modulate antibody pharmacokinetics, a two compartment PK model was implemented where the depot was considered as the first compartment (Figure 6d, Figure 6e). ^47,62^ Using the elimination and absorption rate constants determined from the IV and SC bolus data, the half-life of Centi-C10 from the SC depot compartment, t_1/2_,_dpt_, was found to be 6.7 ± 1.9 hrs from the model. Utilizing the Ritger-Peppas drug release model, time to 50% release can be estimated to be t_50%_ = 32 hrs based on the antibody diffusivity values measured by FRAP and assuming a spherical depot of the same administered volume (Figure S8).^51^ The depot half-life determined from the compartment model of the PK data is significantly smaller than expected from in vitro diffusivity data. The rate constant describing antibody release from the depot, k_dpt_, determined from the model captures all processes leading to loss of antibody mass from the depot with no granularity. It is known that antibodies may be cleared from the subcutaneous space by both active and passive transport mechanisms.^12^ We anticipated that the dominant mechanism of antibody transport out of the hydrogel depot would be passive diffusion; however, given that the depot half-life is shorter than expected, we hypothesize that cells could be infiltrating into the subcutaneous hydrogel depot and causing loss of antibody drug mass by actively trafficking the antibodies or degrading them through consumption or production of reactive oxygen species, or other possible processes. Previous work has shown that immune cells can infiltrate into these PNP hydrogel materials when certain types of cargo such as vaccine components are present and that encapsulated therapeutic cells exhibit motility, indicating that an active antibody transport mechanism could be possible.^31,34^ It is also possible that biological interactions with the material change its physical properties over time, which could alter its drug release characteristics. ^63^ Future work should include investigating cellular infiltration into antibody-loaded PNP hydrogels as well as measuring antibody biodistribution beyond the bloodstream, such as the lymphatic system. Understanding whether cellular infiltration is a problem in this context will help guide the design of future biomaterials.

Additionally, a major concern for the subcutaneous delivery of monoclonal antibodies is bioavailability, which can vary but is generally 50-90%.^12^ The bioavailability of Centi-C10 for the subcutaneously administered formulations was quantified from the area under the PK curve (AUC) through day 42 of the study (Figure 7a). Bioavailability from the SC bolus was 92 ± 11% and the SC gel approached 100% (Figure 7b). Another facet of bioavailability is whether the antibodies will experience degradation and loss of therapeutic function, in this case neutralizing activity, prior to reaching circulation, even if their binding activity remains intact. At Day 7 the neutralizing activity of the antibodies present in the serum was tested in a spike-pseudotyped lentivirus neutralization assay (Figure 7a). There was no difference between the three methods of administration; IV bolus, SC bolus, and SC gel all demonstrated neutralizing activity of 74 ± 12%, 75 ± 16%, and 87 ± 7%, respectively, at a 1:50 dilution in the assay (Figure 7c).

**Figure 7:**
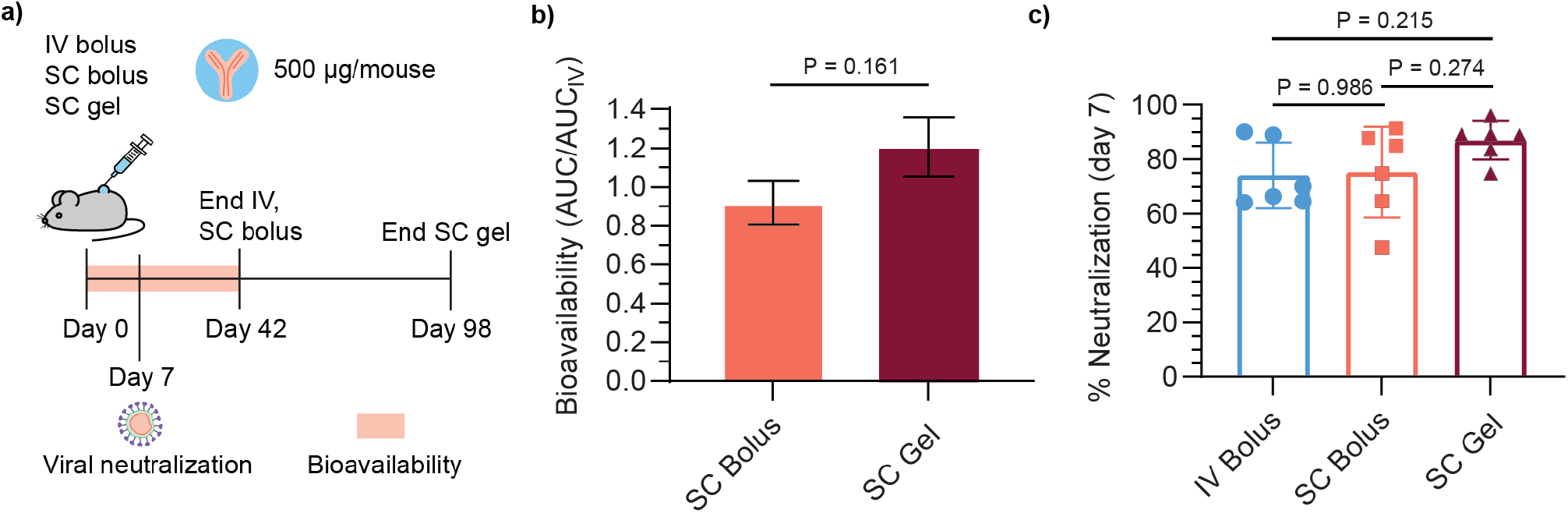
Bioavailability and neutralizing activity of subcutaneously delivered Centi-C10. a) Timecourse of pharmacokinetic study. b) Bioavailability quantified by the area under the curve (AUC) of the Centi-C10 pharmacokinetic profiles by ELISA through day 42 normalized by AUC of the IV bolus group (mean ± SE, n=6 mice/group); p-value determined by two-sided unpaired t-test. c) Neutralizing activity of Centi-C10 from mouse serum 7 days post-administration as determined by spike-pseudotyped lentivirus neutralization assay (mean ± SD, n=6 mice/group, assayed in duplicate). Statistical significance was determined by one-way ANOVA (F (DFn, DFd) = F (2, 15) = 1.906). Tukey post-hoc tests were applied to account for multiple comparisons.

Similar pharmacokinetics were observed for Centi-C10 delivered in a subcutaneous hydrogel formulation in a histidine/sucrose buffer, indicating that buffer exchange is feasible in this system (Figure S5). In addition, biocompatibility was assessed by histology samples of skin surrounding the subcutaneous hydrogel depots (Figure S6, Figure S7). Previous work has shown that PNP hydrogels are biocompatible and are cleared from the subcutaneous space within several weeks, depending on the formulation and injection volume.^31,50^

### 3.5 Extrapolating to human pharmacokinetics and design requirements for depot materials

Beyond evaluating drug depot performance, simple pharmacokinetic modeling can be an important tool to aid the biomaterials design process. In particular, modeling can help inform how a given materials platform, such as the PNP hydrogel system, could be tailored to deliver biotherapeutics such as antibodies with similar physical characteristics but different pharmacokinetic characteristics. Because pharmacokinetics in a mouse model are different than in humans due to differences in physiology and allometric scaling, basic compartment modeling can be used to estimate PK profiles in humans based on both drug and material depot characteristics. ^64^ In order to extrapolate PK profiles in humans, we implemented a two compartment model with first order kinetics and relevant dosing and physiological values for IgG antibodies in humans (Figure 8a). ^12,15,62,62,65^ We used this model to simulate PK profiles for four antibody drugs with distinct elimination half-lives that are either on the market or in preclinical or clinical development (Figure 8b-e). The drugs selected were 1) IgMs and IgG3s, which are a both promising antibody drug subtypes hindered by poor pharmacokinetics (t_1/2,elim_ = 7 days (endogenous)), 2) VRC01, a bnAb against HIV (t_1/2_,_elim_ = 14 days), 3) REGN-COV2, Regeneron’s dual antibody cocktail against SARS-CoV-2 (t_1/2_,_elim_ = 27 days), and 4) mAb-LS, an LS-modified anti-HIV mAb with representative extended half-life characteristics (t,_elim_= 50 days’).^1,13,66–70^ For each drug, profiles were simulated for no depot (single compartment model) compared to a two compartment model with varying depot release half-lives from the material depot compartment to the subcutaneous space. In addition, the time above a threshold of 3 *μ*g/mL serum concentration was quantified, as two of the four selected drugs are representative anti-HIV antibodies, and a threshold concentration of 3 *μ*/mL has been suggested in a simian HIV model as a protective serum titer for passive immunization.^13^ The threshold was kept consistent for sake of comparison.

**Figure 8:**
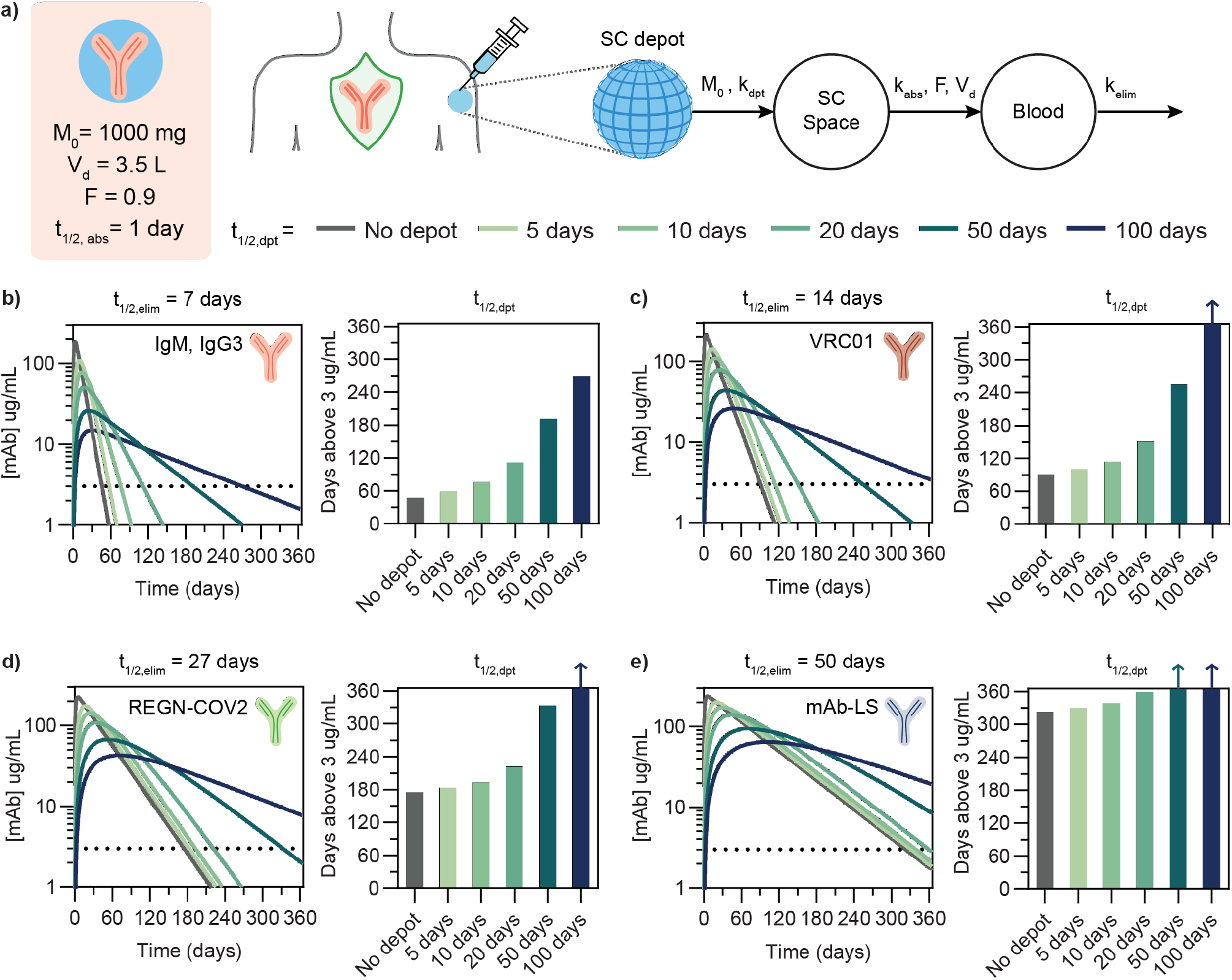
Compartment modeling to understand depot pharmacokinetics in humans. a) two compartment model schema with first-order kinetics between compartments, including relevant physiological parameters and antibody drug characteristics. Pharmacokinetic profiles were generated for antibodies with different elimination half-lives to simulate antibody release from subcutaneous depots exhibiting different release rate constants: b) IgM class antibodies, c) VRC01, an anti-HIV bnAb, d) REGN-COV2, antibody cocktail against SARS-CoV-2, and e) mAb-LS, indicating a bnAb modified with the LS mutation to increase serum half-life. Arrows on bars indicate that serum titer remained above the 3 *μ*g/mL threshold beyond 365 days.

For antibodies such as IgM or IgG3 with a relatively short elimination half-life of 7 days, a drug depot with a release half-life of 10 days increases the duration above the therapeutic threshold by 1.6 fold and a depot halflife of 20 days yields a 2.4 fold duration increase. For the anti-HIV bnAb VRC01 (t_1/2,elim_ = 14 days), a depot half-life of 20 days increases duration by 1.7 fold. For a neutralizing antibody cocktail like REGN-COV2 with a comparatively longer half-life of 27 days, a depot half-life of 50 days would be required to extend the duration above threshold by 2 fold. As drug elimination half-life increases, the elimination phase becomes the primary determinant of the overall PK profile; in the case of a long-lived antibody such as an LS mAb with t_1/2,elim_ = 50 days, the depot would need to exhibit very slow release to modify the PK profile. In order for the depot compartment to meaningfully extended PK profiles, the depot release half-life needs to significantly longer than the elimination half life so it becomes the rate limiting step. From the compartment modeling of these representative antibody drugs, a general trend emerges that in order for the duration above threshold to increase 2 fold, the depot half-life must be approximately 2 fold greater than the drug elimination half life. Depending on the drug characteristics and desired PK profile for its target application, this implies that antibodies with poor PK characteristics may benefit the most from depot formulation technology. For a 2:10 hydrogel formulation administered subcutaneously at a suitable volume for humans (1.5 mL), we can estimate time to 50% release as t_50%_ = 6.3 days based on in vitro antibody diffusivity measurements.^51,71^ The current formulation could be more effective for short-lived antibody drugs.

Employing a relatively simple two compartment model is useful not only for estimating antibody delivery from a subcutaneous depot, but it can also be adapted and applied to other delivery platforms in which the benchmark for success is the ability to maintain serum drug concentration either above a threshold or within a therapeutic window over time. We acknowledge that much more sophisticated pharmacokinetic and pharmacodynamic models exist; however, in the case of antibodies, a simple model is generally sufficient to be used as a benchmarking tool to estimate the observed PK by capturing the dominant mass transport processes in vivo. Incorporating compartment modeling during the depot design process is important to estimate what depot release kinetics are required to achieve a desired PK profile and ensure that the depot design is relevant for translation.

## 4 Conclusions

PNP hydrogels exhibit many of the key attributes necessary for a subcutaneous delivery depot, including injectability, prolonged drug exposure, and maintenance of drug stability during stressed aging. An in vivo PK study comparing delivery of a SARS-Cov-2 antibody via bolus administration and encapsulated in a PNP hydrogel showed that the hydrogel was able to deliver functional antibody, but that the release kinetics from the depot were faster than expected based on in vitro experiments. This work highlights both the future potential and reiterates outstanding challenges of delivering antibody therapeutics from a subcutaneous depot. Implementing a two compartment pharmacokinetic model to simulate the required depot drug kinetics for clinical or preclinical human antibodies can inform the design of new materials that could serve as subcutaneous biotherapeutic delivery depots for applications in infectious diseases including pre- and post-exposure prophylaxis, treatment, and passive immunization, as well as other applications in biotherapeutic delivery.

## Author Contributions

C.M.K., A.C.Y., A.E.P., G.A.R., C.K.J., P.S.K., S.Y., J.G., and E.A.A. designed experiments. C.M.K., A.C.Y., A.E.P., C.K.J. and A.B. conducted experiments. C.S.L. assisted with animal studies. C.M.K., A.C.Y., A.E.P., C.K.J., C.L.M. and E.A.A. analyzed data. C.M.K., A.C.Y., A.E.P., C.L.M., S.Y., J.G., and E.A.A. wrote the manuscript.

## Conflicts of interest

Centivax has patented Centi-10, as it may have commercial relevance to the field of virology.

## Acknowledgements

This research was financially supported by the Center for Human Systems Immunology with the Bill & Melinda Gates Foundation (OPP1113682; OPP1211043) and the National Institutes of Health (NIAID R01 AI154989). C.M.K. was supported by a Stanford Graduate Fellowship and the Stanford Bio-X William and Lynda Steere Fellowship. A.C.Y. was supported by a Kodak Fellowship. A.E.P. was supported by the Stanford Maternal and Child Health Research Institute postdoctoral fellowship. C.S.L. was supported by a Department of Defense National Defense Science and Engineering Graduate (NDSEG) Fellowship. C.K.J. was supported by a National Science Foundation (NSF) Graduate Research Fellowship. C.L.M. was supported by the NSERC Postgraduate Scholarship and the Stanford BioX Bowes Graduate Student Fellowship. The authors also thank every member of the Appel Lab, former and current, for their on-going support, technical expertise, and scientific discussion. In particular, the authors thank Abigail K. Grosskopf, Olivia M. Saouaf, Andrea D’Aquino and Noah Eckman for their technical assistance with rheometry, FRAP, polymer synthesis, and MATLAB, respectively. FRAP measurements and histology images were acquired at the Stanford University Cell Sciences Imaging Core Facility (CSIF, RRID: SCR_017787) on equipment funded jointly by the Stanford School of Engineering and the Beckman Center. The authors also thank Eric E. Petersen and Doreen Wu in the Stanford Animal Histology Services for help with preparation of histologic specimens.

## Data Availability

All data supporting the results in this study are available within the article and its Supplementary Information. Any raw datasets acquired and analyzed (or any subsets thereof), which would require contextual metadata for reuse, are available from the corresponding author upon reasonable request.

## Notes

### Competing Interest Statement

Centivax has patented Centi-C10, as it may have commercial relevance to the field of virology.

## References

[1] O’Brien, M. P.; Forleo-Neto, E.; Musser, B. J.; Isa, F.; Chan, K.-C.; Sarkar, N.; Bar, K. J.; Barnabas, R. V.; Barouch, D. H.; Cohen, M. S., et al. New England Journal of Medicine 2021, 385, 1184–1195.

[2] Abraham, J. Nature Reviews Immunology 2020, 20, 401–403.

[3] Burton, D. R. Nature Reviews Immunology 2019, 19, 77–78.

[4] Morris, L.; Mkhize, N. N. PLoS Medicine 2017, 14, e1002436.

[5] Cockburn, I. A.; Seder, R. A. Nature immunology 2018, 19, 1199–1211.

[6] Barrett, A. D. npj Vaccines 2018, 3, 1–4.

[7] Hooft van Huijsduijnen, R.; Kojima, S.; Carter, D.; Okabe, H.; Sato, A.; Akahata, W.; Wells, T. N.; Katsuno, K. PLoS Neglected Tropical Diseases 2020, 14, e0007860.

[8] Domachowske, J. Vaccines; Springer, 2021; pp 13–23.

[9] Ryman, J. T.; Meibohm, B. CPT: pharmacometrics & systems pharmacology 2017, 6, 576–588.

[10] Mitragotri, S.; Burke, P. A.; Langer, R. Nature reviews Drug discovery 2014, 13, 655–672.

[11] Caskey, M.; Klein, F.; Nussenzweig, M. C. Nature medicine 2019, 25, 547–553.

[12] Viola, M.; Sequeira, J.; Seiça, R.; Veiga, F.; Serra, J.; Santos, A. C.; Ribeiro, A. J. Journal of controlled release 2018, 286, 301–314.

[13] Gautam, R.; Nishimura, Y.; Gaughan, N.; Gazumyan, A.; Schoofs, T.; Buckler-White, A.; Seaman, M. S.; Swihart, B. J.; Follmann, D. A.; Nussenzweig, M. C., et al. Nature medicine 2018, 24, 610–616.

[14] Mahomed, S.; Garrett, N.; Baxter, C.; Abdool Karim, Q.; Abdool Karim, S. S. The Journal of Infectious Diseases 2021, 223, 370–380.

[15] Mahomed, S.; Garrett, N.; Capparelli, E. V.; Osman, F.; Harkoo, I.; Yende-Zuma, N.; Gengiah, T. N.; Archary, D.; Samsunder, N.; Baxter, C., et al. The Journal of Infectious Diseases 2022,

[16] Wang, W.; Singh, S.; Zeng, D. L.; King, K.; Nema, S. Journal of pharmaceutical sciences 2007, 96, 1–26.

[17] Correa, S.; Grosskopf, A. K.; Lopez Hernandez, H.; Chan, D.; Yu, A. C.; Stapleton, L. M.; Appel, E. A. Chemical Reviews 2021, 121, 11385–11457.

[18] Webber, M. J.; Appel, E. A.; Meijer, E.; Langer, R. Nature materials 2016, 15, 13–26.

[19] Mazutis, L.; Vasiliauskas, R.; Weitz, D. A. Macromolecular bioscience 2015, 15, 1641–1646.

[20] Ferreira, N. N.; Caetano, B. L.; Boni, F. I.; Sousa, F.; Magnani, M.; Sarmento, B.; Cury, B. S. F.; Gremião, M. P. D. Journal of Pharmaceutical Sciences 2019, 108, 1559–1568.

[21] Schweizer, D.; Vostiar, I.; Heier, A.; Serno, T.; Schoenhammer, K.; Jahn, M.; Jones, S.; Piequet, A.; Beerli, C.; Gram, H., et al. Journal of Controlled Release 2013, 172, 975–982.

[22] Schweizer, D.; Schonhammer, K.; Jahn, M.; Gopferich, A. Biomacromolecules 2013, 14, 75–83.

[23] Schweizer, D.; Serno, T.; Goepferich, A. European Journal of Pharmaceutics and Biopharmaceutics 2014, 88, 291–309.

[24] Sousa, F.; Cruz, A.; Fonte, P.; Pinto, I. M.; Neves-Petersen, M. T.; Sarmento, B. Scientific reports 2017, 7, 1–13.

[25] Koutsopoulos, S.; Zhang, S. Journal of controlled release 2012, 160, 451–458.

[26] Lovett, M. L.; Wang, X.; Yucel, T.; York, L.; Keirstead, M.; Haggerty, L.; Kaplan, D. L. European journal of pharmaceutics and biopharmaceutics 2015, 95, 271–278.

[27] Meis, C. M.; Salzman, E. E.; Maikawa, C. L.; Smith, A. A.; Mann, J. L.; Grosskopf, A. K.; Appel, E. A. ACS Biomaterials Science & Engineering 2020,

[28] Appel, E. A.; Tibbitt, M. W.; Webber, M. J.; Mattix, B. A.; Veiseh, O.; Langer, R. Nature communications 2015, 6, 1–9.

[29] Yu, A. C.; Smith, A. A.; Appel, E. A. Molecular Systems Design & Engineering 2020, 5, 401–407.

[30] Stapleton, L. M.; Steele, A. N.; Wang, H.; Hernandez, H. L.; Anthony, C. Y.; Paulsen, M. J.; Smith, A. A.; Roth, G. A.; Thakore, A. D.; Lucian, H. J., et al. Nature biomedical engineering 2019, 3, 611–620.

[31] Roth, G. A.; Gale, E. C.; Alcántara-Hernández, M.; Luo, W.; Axpe, E.; Verma, R.; Yin, Q.; Yu, A. C.; Lopez Hernandez, H.; Maikawa, C. L., et al. ACS central science 2020, 6, 1800–1812.

[32] Lopez Hernandez, H.; Grosskopf, A. K.; Stapleton, L. M.; Agmon, G.; Appel, E. A. Macromolecular bioscience 2019, 19, 1800275.

[33] Grosskopf, A. K.; Roth, G. A.; Smith, A. A.; Gale, E. C.; Hernandez, H. L.; Appel, E. A. Bioengineering & translational medicine 2020, 5, e10147.

[34] Grosskopf, A. K.; Labanieh, L.; Klysz, D. D.; Roth, G.; Xu, P.; Adebowale, O.; Gale, E. C. G. C.; Jons, C. K.; Klich, J. H.; Yan, J., et al. bioRxiv 2021,

[35] Correa, S.; Gale, E. C.; Mayer, A. T.; Xiao, Z.; Liong, C.; Klich, J. H.; Brown, R. A.; Meany, E.; Saouaf, O.; Maikawa, C. L., et al. bioRxiv 2021,

[36] Gale, E. C.; Lahey, L. J.; Böhnert, V; Powell, A. E.; Ou, B. S.; Carozza, J. A.; Li, L.; Appel, E. A. bioRxiv 2021,

[37] Gale, E. C.; Powell, A. E.; Roth, G. A.; Meany, E. L.; Yan, J.; Ou, B. S.; Grosskopf, A. K.; Adamska, J.; Picece, V. C.; d’Aquino, A. I., et al. Advanced Materials 2021, 33, 2104362.

[38] Grosskopf, A. K.; Correa, S.; Baillet, J.; Maikawa, C. L.; Gale, E. C.; Brown, R. A.; Appel, E. A. Communications biology 2021, 4, 1–7.

[39] Roth, G. A.; Saouaf, O. M.; Smith, A. A.; Gale, E. C.; Hernandez, M. A.; Idoyaga, J.; Appel, E. A. ACS biomaterials science & engineering 2021, 7, 1889–1899.

[40] Saouaf, O. M.; Roth, G. A.; Ou, B. S.; Smith, A. A.; Yu, A. C.; Gale, E. C.; Grosskopf, A. K.; Picece, V. C.; Appel, E. A. Journal of Biomedical Materials Research Part A 2021, 109, 2173–2186.

[41] Stapleton, L. M.; Lucian, H. J.; Grosskopf, A. K.; Smith, A. A.; Totherow, K. P.; Woo, Y. J.; Appel, E. A. Advanced Therapeutics 2021, 4, 2000242.

[42] Meis, C. M.; Grosskopf, A. K.; Correa, S.; Appel, E. A. JoVE (Journal of Visualized Experiments) 2021, e62234.

[43] Axelrod, D.; Koppel, D.; Schlessinger, J.; Elson, E.; Webb, W. W. Biophysical journal 1976, 16, 1055–1069.

[44] Powell, A. E.; Zhang, K.; Sanyal, M.; Tang, S.; Weidenbacher, P. A.; Li, S.; Pham, T. D.; Pak, J. E.; Chiu, W.; Kim, P. S. ACS central science 2021, 7, 183–199.

[45] Crawford, K. H.; Eguia, R.; Dingens, A. S.; Loes, A. N.; Malone, K. D.; Wolf, C. R.; Chu, H. Y.; Tortorici, M. A.; Veesler, D.; Murphy, M., et al. Viruses 2020, 12, 513.

[46] Rogers, T. F.; Zhao, F.; Huang, D.; Beutler, N.; Burns, A.; He, W.-T.; Limbo, O.; Smith, C.; Song, G.; Woehl, J., et al. Science 2020, 369, 956–963.

[47] Zou, H.; Banerjee, P.; Leung, S. S. Y.; Yan, X. Frontiers in Pharmacology 2020, 11, 997.

[48] Lopez Hernandez, H.; Souza, J. W.; Appel, E. A. Macromolecular Bioscience 2021, 21, 2000295.

[49] Grosskopf, A. K.; Saouaf, O. A.; Lopez Hernandez, H.; Appel, E. A. Journal of Polymer Science 2021, 59, 2854–2866.

[50] Jons, C. K.; Grosskopf, A. K.; Baillet, J.; Yan, J.; Klich, J. H.; Appel, E. A. bioRxiv 2022,

[51] Ritger, P. L.; Peppas, N. A. Journal of controlled release 1987, 5, 37–42.

[52] Jøssang, T.; Feder, J.; Rosenqvist, E. Journal of protein chemistry 1988, 7, 165–171.

[53] Armstrong, J. K.; Wenby, R. B.; Meiselman, H. J.; Fisher, T. C. Biophysical journal 2004, 87, 4259–4270.

[54] Chukwu, O. A.; Adibe, M. The International journal of health planning and management 2022, 37, 930–943.

[55] Songane, M. International Journal of Health Economics and Management 2018, 18, 197–219.

[56] Controlled Temperature Chain Working Group, Controlled temperature chain: strategic roadmap for priority vaccines 2017-2020, World Health Organization 2017(WHO/IVB/17.20.

[57] Chaudhury, C.; Mehnaz, S.; Robinson, J. M.; Hayton, W. L.; Pearl, D. K.; Roopenian, D. C.; Anderson, C. L. Journal of Experimental Medicine 2003, 197, 315–322.

[58] Roopenian, D. C.; Christianson, G. J.; Sproule, T. J.; Brown, A. C.; Akilesh, S.; Jung, N.; Petkova, S.; Avanessian, L.; Choi, E. Y.; Shaffer, D. J., et al. The Journal of Immunology 2003, 170, 3528–3533.

[59] Keizer, R. J.; Huitema, A. D.; Schellens, J. H.; Beijnen, J. H. Clinical pharmacokinetics 2010, 49, 493–507.

[60] Ko, S.; Jo, M.; Jung, S. T. BioDrugs 2021, 35, 147–157.

[61] Proetzel, G.; Roopenian, D. C. Methods 2014, 65, 148–153.

[62] Kagan, L. Drug Metabolism and Disposition 2014, 42, 1890–1905.

[63] Chen, W.; Yung, B. C.; Qian, Z.; Chen, X. Advanced drug delivery reviews 2018, 127, 20–34.

[64] Nair, A. B.; Jacob, S. Journal of basic and clinical pharmacy 2016, 7, 27.

[65] Stephenson, K. E.; Julg, B.; Tan, C. S.; Zash, R.; Walsh, S. R.; Rolle, C.-P.; Monczor, A. N.; Lupo, S.; Gelderblom, H. C.; Ansel, J. L., et al. Nature medicine 2021, 27, 1718–1724.

[66] Mayer, K. H.; Seaton, K. E.; Huang, Y.; Grunenberg, N.; Isaacs, A.; Allen, M.; Ledgerwood, J. E.; Frank, I.; Sobieszczyk, M. E.; Baden, L. R., et al. PLoS medicine 2017, 14, e1002435.

[67] Gautam, R.; Nishimura, Y.; Pegu, A.; Nason, M. C.; Klein, F.; Gazumyan, A.; Golijanin, J.; Buckler-White, A.; Sadjadpour, R.; Wang, K., et al. Nature 2016, 533, 105–109.

[68] Ko, S.-Y.; Pegu, A.; Rudicell, R. S.; Yang, Z.-y.; Joyce, M. G.; Chen, X.; Wang, K.; Bao, S.; Kraemer, T. D.; Rath, T., et al. Nature 2014, 514, 642–645.

[69] Chu, T. H.; Patz Jr, E. F.; Ackerman, M. E. Coming together at the hinges: therapeutic prospects of IgG3. MAbs. 2021; p 1882028.

[70] Keyt, B. A.; Baliga, R.; Sinclair, A. M.; Carroll, S. F.; Peterson, M. S. Antibodies 2020, 9, 53.

[71] Badkar, A. V.; Gandhi, R. B.; Davis, S. P.; LaBarre, M. J. Drug Design, Development and Therapy 2021, 15, 159.

