## Supplemental Information for "Subcutaneous delivery of an antibody against SARS-CoV-2 from a supramolecular hydrogel depot"

### Contents

**1 Supplemental Methods** **S3**
1.1 Formulation and in vivo pharmacokinetic study of Centi-C10 antibody in histidine buffer . . . S3
1.2 Histology sample preparation . . . S3
1.3 Pharmacokinetic modeling . . . S3
1.3.1 One compartment model . . . S3
1.3.2 Two compartment model . . . S3

**2 Centi-C10 antibody sequence** **S5**

**3 Additional PNP hydrogel rheology** **S6**
3.1 Rheological stability . . . S6
3.2 Viscosity recovery time from step shear test . . . S6

**4 Fluorescence recovery after photobleaching (FRAP) data** **S7**

**5 In vitro Centi-C10 stability at 50 °C** **S7**

**6 In vivo PK data for Centi-C10 antibody in hydrogel with histidine buffer** **S8**

**7 Histology - Day 7** **S9**

**8 Modeling depot drug release as a function of cargo diffusivity** **S10**

### 1 Supplemental Methods

#### 1.1 Formulation and in vivo pharmacokinetic study of Centi-C10 antibody in histidine buffer

Prior to hydrogel formulation, the Centi-C10 antibody was buffer exchanged from PBS into 25 mM histidine buffer (pH 6) (sterile, Bioworld) with 150 mM USP-grade sucrose (Sigma-Aldrich) using spin desalting columns. Hydrogel stock solution components were prepared in the same histidine buffer, and the hydrogels were formulated as described in the paper. The in vivo pharmacokinetic study was conducted as described in the paper.

#### 1.2 Histology sample preparation

A parallel cohort of mice (n=3) to the PK study were subcutaneously administered both unloaded hydrogel and hydrogel loaded with Centi-C10 antibody on separate flanks. At day 7, the mice were euthanized and the hydrogel and surrounding subcutaneous tissue were excised, embedded in OCT (optimal cutting temperature) medium in block molds, and flash frozen with liquid nitrogen. Frozen samples were submitted to Stanford Animal Histology Services for sectioning, mounting, and standard staining.

#### 1.3 Pharmacokinetic modeling

##### 1.3.1 One compartment model

The differential equations and the analytical solution for a standard one compartment pharmacokinetic model with first order reaction kinetics are described in Zou, et al.[1] Briefly, the form of the analytical solution of the single compartment model for drug serum concentration as a function of time, *C(t)*, used in our modeling is as follows:

$$C(t) = \frac{F k_{abs} M_0}{V_d (k_{abs} - k_{elim})} \left( e^{-k_{elim} t} - e^{-k_{abs} t} \right) \tag{S1}$$

where $F$ = bioavailability, $k_{abs}$ = rate constant of absorbance from subcutaneous space into the bloodstream, $M_0$ = initial dose of drug, $V_d$ = volume of distribution, $k_{elim}$ = rate constant of drug elimination, and $t$ = time.

##### 1.3.2 Two compartment model

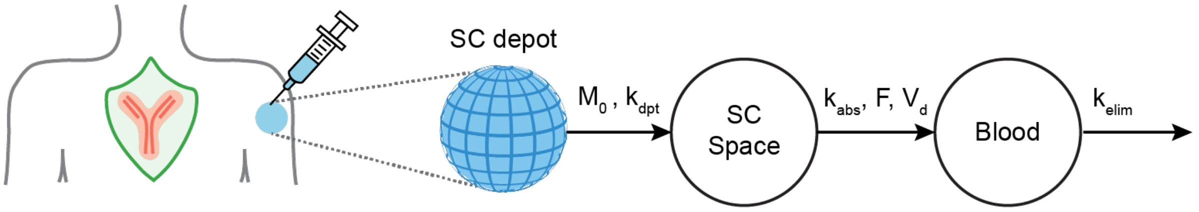


**Figure S1:** Two compartment model schema

The following differential equations and initial conditions were written to describe the mass transport depicted in Figure S1, where $M_1$ refers to drug mass in the subcutaneous hydrogel depot compartment, $M_2$ refers to drug mass in the subcutaneous compartment, and $M_3$ refers drug mass in the blood serum compartment:

$$\frac{dM_1(t)}{dt} = -k_{dpt} M_1(t), \quad M_1(0) = M_0 \tag{S2}$$

$$\frac{dM_2(t)}{dt} = k_{dpt} M_1(t) - k_{abs} M_2(t), \quad M_2(0) = 0 \tag{S3}$$

$$\frac{dM_3(t)}{dt} = k_{abs}FM_2(t) - k_{elim}M_3(t), \quad M_3(0) = 0 \tag{S4}$$

where $k_{dpt}$ = rate constant of drug release from hydrogel depot, and

$$C(t) = \frac{M_3(t)}{V_d} \tag{S5}$$

These equations and conditions were solved for the following analytical solution:

$$C(t) = \frac{M_0 F k_{dpt} k_{abs} e^{-k_{elim}t}}{V_d(k_{dpt} - k_{abs})(k_{elim} - k_{dpt})(k_{elim} - k_{abs})} \left[ (k_{elim} - k_{dpt}) e^{(k_{elim} - k_{abs})t} + (k_{abs} - k_{elim}) e^{(k_{elim} - k_{dpt})t} + k_{dpt} - k_{abs} \right] \tag{S6}$$

#### 2 Centi-C10 antibody sequence

**VH**
QVQLVQSGAEVKKPGSSVKVSCKASGYPFTNYGISWVRQAPGQGLEWMGWMNPNSGNTGYAQKFQ
GRVTITADESTSTAYMELSSLRSEDTAVYYCATLSGISTPMDVWGQGTLVTVSS

**VL**
DIVMTQSPDSLAVSLGERATINCRSSQSVLYSSNNKNYFAWYQQKPGQPPKLLIYWASIRGSGVPDRFSGS
GSGTDFTLTISSLQAEDVAVYYCHQYYTTPPTFGGGTKVEIKR

### 3 Additional PNP hydrogel rheology

#### 3.1 Rheological stability

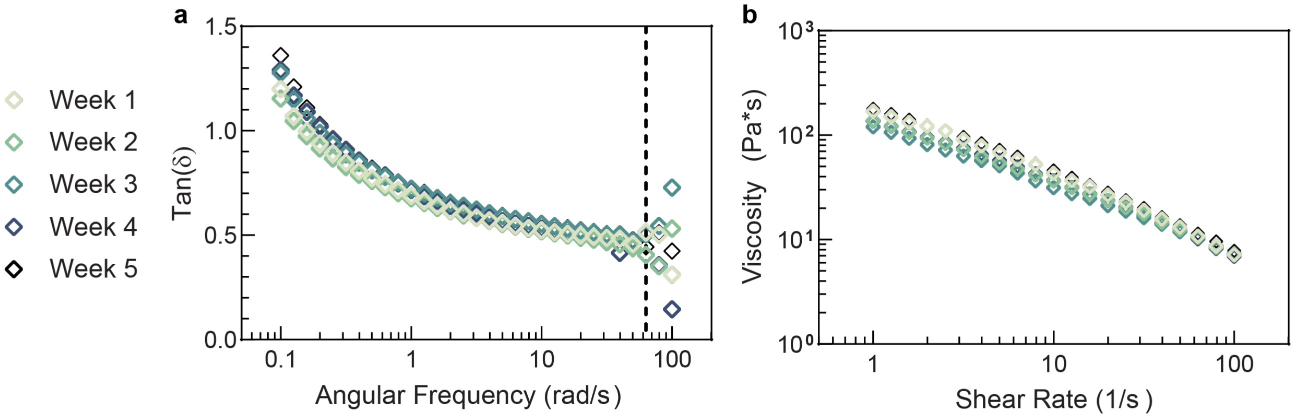


**Figure S 2:** Relative elasticity and steady shear measurements. (a) Tan($\delta$) for the high concentration gel measured once every week for 5 weeks illustrates minimal change in relative elasticity. The dotted line indicates the angular frequency cutoff, where higher values are likely influenced by inertial effects. (b) Steady shear measurements of the high concentration gel measured once every week for 5 weeks illustrates minimal change in viscosity or shear-thinning index.

#### 3.2 Viscosity recovery time from step shear test

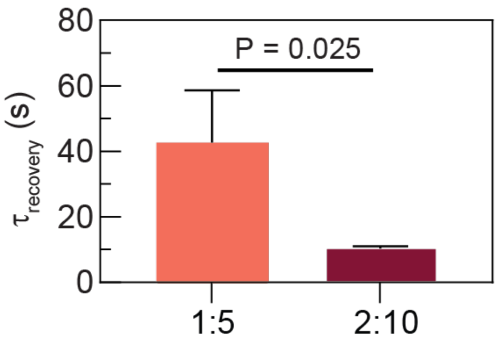


**Figure S3:** Characteristic recovery time $\tau$ was determined from the viscosity recovery curves from the step shear tests shown in Figure 2d (mean ± SD, n=3, final three cycles); p-value determined by two-sided, unpaired t-test.

#### 4 Fluorescence recovery after photobleaching (FRAP) data

**Table S1.** Diffusivity values as determined by FRAP. Mean ±SD (n=3)

| Gel formulation | $D_{IgG}$ ($\mu m^2/s$) | $D_{polymer}$ ($\mu m^2/s$) | $D_{IgG}/D_{polymer}$ |
|---|---|---|---|
| 1:5 | 4.3 ±0.4 | 1.3 ±0.2 | 3.4 ±0.7 |
| 2:10 | 0.9 ±0.2 | 1.4 ±0.1 | 0.6 ±0.1 |

#### 5 In vitro Centi-C10 stability at 50 °C

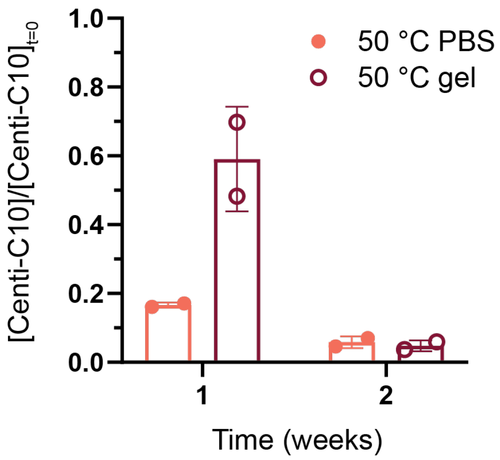


**Figure S4:** In vitro stability of Centi-C10 in PBS buffer formulation compared to hydrogel encapsulated Centi-C10 at 50 °C quantified via anti-RBD ELISA (mean ± SD, assayed in duplicate)

### 6 In vivo PK data for Centi-C10 antibody in hydrogel with histidine buffer

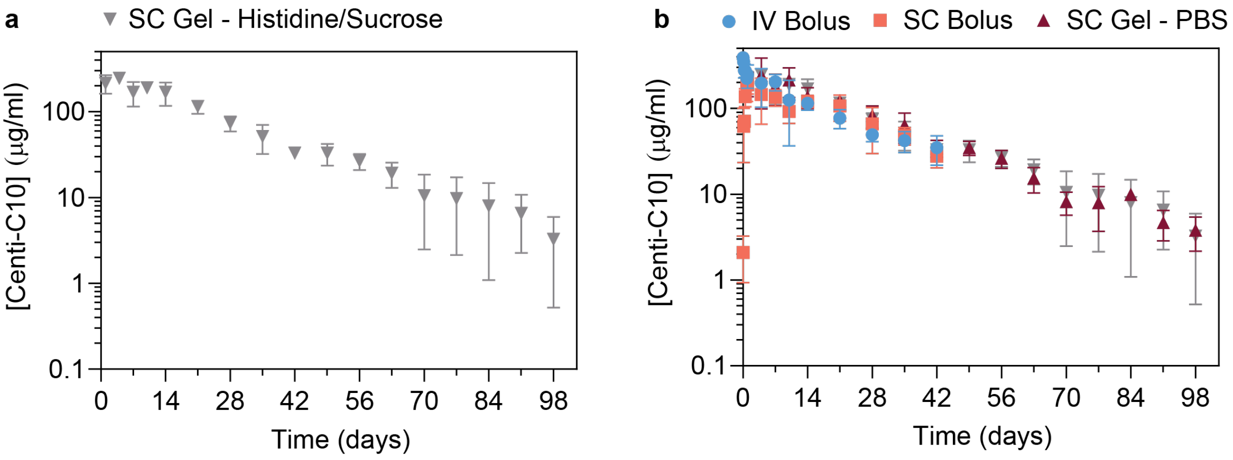


**Figure S5:** Centi-C10 serum pharmacokinetics in a preclinical mouse model as determined by ELISA. a) PK profile for Centi-C10 administered from a SC gel with histidine/sucrose buffer. b) SC gel with histidine/sucrose buffer overlaid with IV, SC bolus, and SC gel with PBS as shown in the paper. Data points shown as mean ± SD (n = 3 prior to 24 hrs, otherwise n = 6; for gel groups, n ≥ 4 after day 63).

#### 7 Histology - Day 7

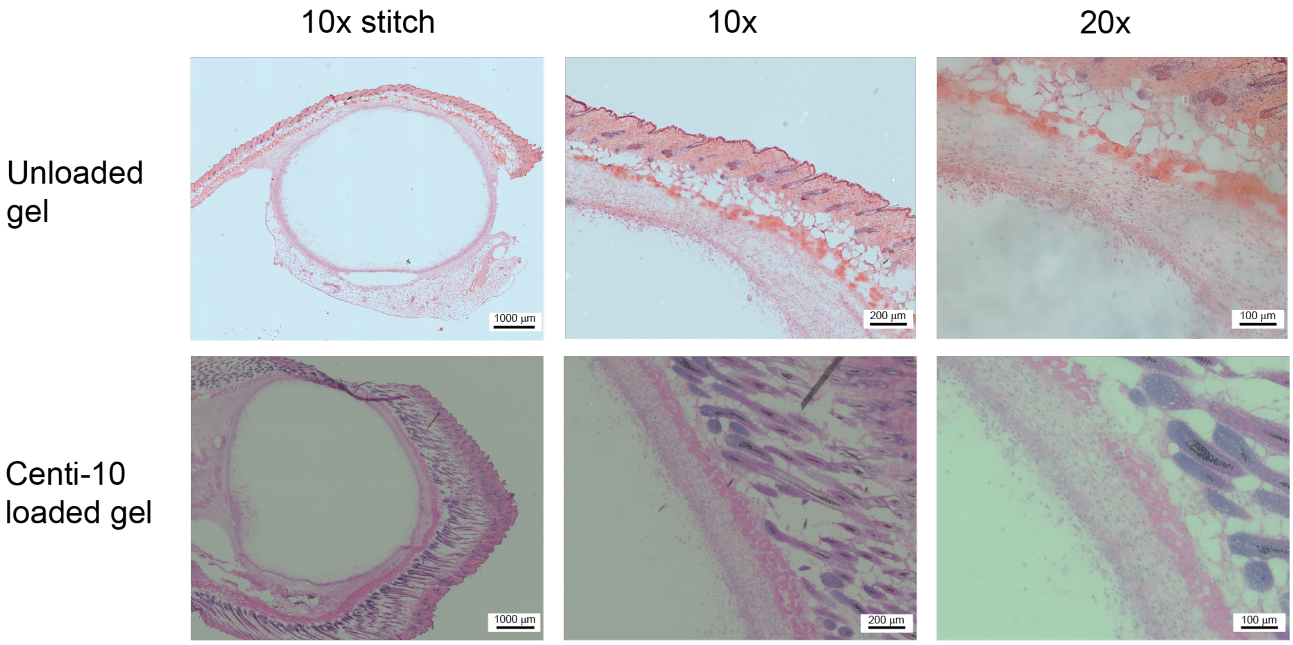


**Figure S6:** Hematoxylin and eosin (H E) staining, representative images at Day 7. Stitched image show entire cross section of excised hydrogel along with surrounding skin tissue.

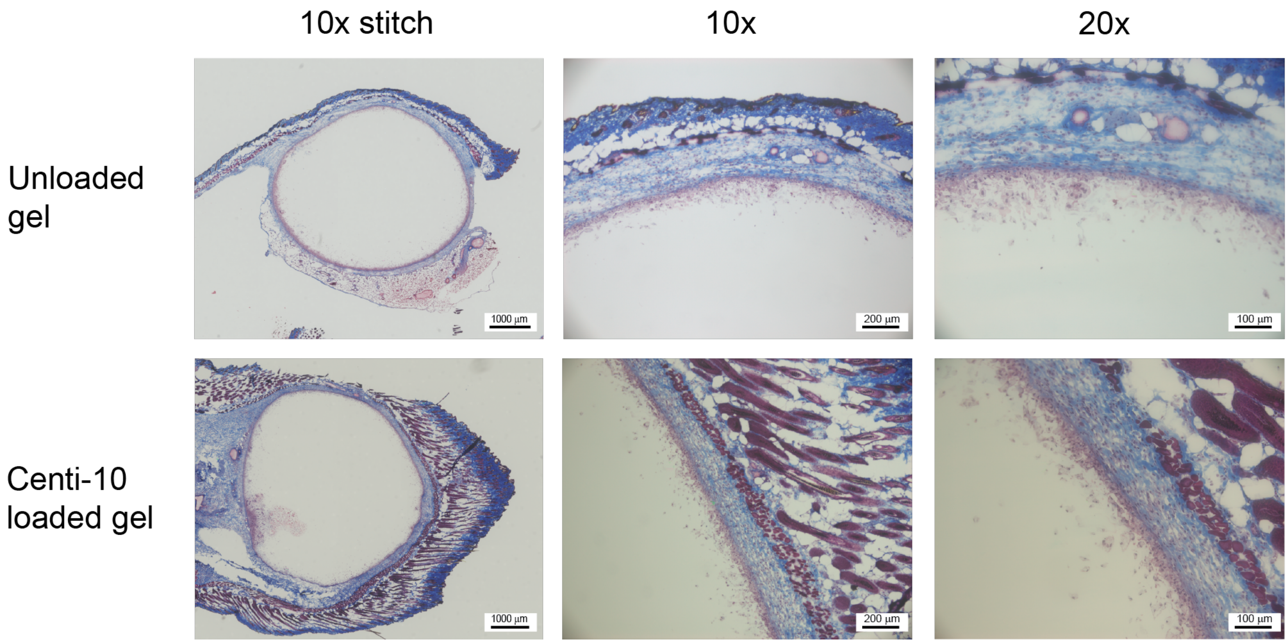


**Figure S7:** Trichrome staining, representative images at Day 7. Stitched image show entire cross section of excised hydrogel along with surrounding skin tissue.

### 8 Modeling depot drug release as a function of cargo diffusivity

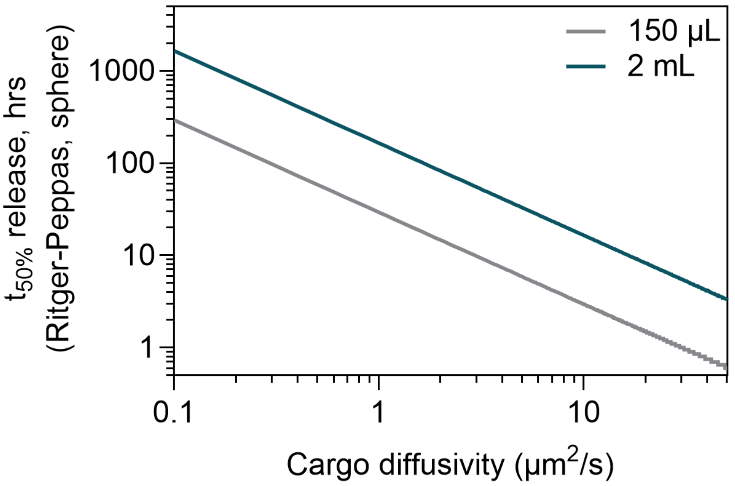


**Figure S8:** Time to 50% release from a depot as a function of encapsulated cargo diffusivity from a 150 $\mu$L depot (mouse) and a 2 mL depot (human), which are typical volumes for subcutaneous injection. Profiles generated in Matlab using the Ritger-Peppas mass release approximation assuming Fickian release and spherical depot.[2]

#### References


[1] Zou, H.; Banerjee, P.; Leung, S. S. Y.; Yan, X. *Frontiers in Pharmacology* **2020**, *11*, 997.

[2] Ritger, P. L.; Peppas, N. A. *Journal of controlled release* **1987**, *5*, 37–42.
